# Mouse model of atypical DAT deficiency syndrome uncovers dopamine dysfunction associated with parkinsonism and psychiatric disease

**DOI:** 10.1101/2023.08.17.553695

**Authors:** Freja Herborg, Lisa K. Konrad, Ciara Pugh, Benoît Delignat-Lavaud, Cecilia Friis Ratner, Nora Awadallah, Jose A. Pino, Frida Berlin, Mathias Rickhag, Birgitte Holst, David Woldbye, Gonzalo Torres, Louis-Eric Trudeau, Ulrik Gether

**Affiliations:** Molecular Neuropharmacology and Genetics Laboratory, Department of Neuroscience, Faculty of Health and Medical Sciences, University of Copenhagen, Copenhagen, Denmark; CNS Research Group, Department of Pharmacology and Physiology, Department of Neurosciences, Faculty of Medicine, Université de Montréal, Montreal, QC, Canada; Department of Biomedical Sciences, University of Copenhagen, Copenhagen, Denmark; Department of Molecular, Cellular, and Biomedical Sciences, City University of New York School of Medicine at City College, New York, United States; Department of Medicine, School of Medicine, University of Atacama, Copiapó, Chile; Department of Molecular Pharmacology and Neuroscience, Stritch School of Medicine, Loyola University, Chicago, United States

## Abstract

The dopamine transporter (DAT) plays a crucial role in regulating the brain’s dopamine (DA) homeostasis. Atypical DAT deficiency syndrome (DTSD) is a disease characterized by early-onset parkinsonism and comorbid psychiatric symptoms, but the pathobiological processes that link DAT dysfunction to both parkinsonism and psychiatric symptoms are unknown. Here, we present a genetic mouse model of atypical DTDS that expresses two coding DAT variants, DAT-I312F and DAT-D421N, derived from a patient diagnosed with ADHD and parkinsonism. Phenotypic characterization of the mutant mice revealed impaired DAT function and major homeostatic changes including increased ambient extracellular DA levels, decreased evoked DA release, and reduced expression of both tyrosine hydroxylase (TH) and of DA D1/D2 receptors. This was accompanied by diminished striatal dopaminergic axonal density and a psychomotor phenotype characterized by hyperactivity, enhanced exploratory activity, and pronounced clasping. Importantly, both amphetamine and anticholinergic treatment ameliorated aberrant hyperlocomotion in the mice. Summarized, by replicating core aspects of the patient’s phenotype, the mouse model not only provides insights into the mechanisms underlying atypical DTDS but also underlines the broad relevance of DA deficits for understanding the co-morbidity between neuropsychiatric diseases and parkinsonism.

**ONE SENTENCE SUMMARY:** In a new mouse disease model, we explore the behavioral consequences and dopaminergic dysfunction that arise from patient-derived mutations in the dopamine transporter associated with parkinsonism and co-morbid neuropsychiatric disease

## INTRODUCTION

Neuropsychiatric and neurodegenerative diseases of the brain are notoriously difficult to treat, resulting in a large unmet need for developing new treatment strategies. Despite differences in clinical presentation and disease onset, accumulating evidence points to the relevance of investigating putative overlapping pathological mechanisms. For instance, the monoaminergic circuits that show high vulnerability in Parkinson’s disease (PD) (1, 2) have been heavily implicated in the pathophysiology of psychiatric diseases (3-7). There is also evidence that neuropsychiatric disorders and neurodegenerative diseases have mutual risk factors. More than 25% of patients diagnosed with neurodegenerative disorders already have a psychiatric diagnosis, and high rates of psychiatric symptoms have been reported for some genetic forms of PD (8-12). This co-morbidity could reflect shared or overlapping etiological factors or commonalities in the molecular and cellular pathways. These factors highlight the need for disease models with construct validity to investigate pathobiological processes.

Dopamine (DA) is recognized for its dual role in basal locomotor control and in regulating emotional states that are critical for higher order brain functions such as attention, motivation and reinforcement learning. Consistently, DA dysfunction has been implicated in both neurodegenerative diseases, particularly PD, and in neuropsychiatric disorders such as Attention-Deficit/Hyperactivity Disorder (ADHD), bipolar disease, schizophrenia, and substance use disorder (3-6, 13). Indeed, core symptoms of these diseases may be alleviated by therapeutics targeting the DA circuits. However, we still do not fully understand how DA exerts its multiple functions and the nature and progression of DA dysfunction in diseased states. Identifying molecular and neural consequences of DA dysfunction, particularly in relation to behavioral phenotypes, may provide critical insights into disease pathology, and thereby support the development of novel pharmacotherapies.

DA homeostasis relies heavily on reuptake via the DA transporter (DAT), which is selectively expressed by DA neurons. DAT thereby ensures a tight spatiotemporal regulation of extracellular DA levels and a synthesis-independent DA supply to dopaminergic terminals (14, 15). The critical function of DAT was underlined when complete loss of function mutations in DAT were shown to cause hereditable infantile parkinsonism dystonia or ‘dopamine transporter deficiency syndromé (DTDS) (16). Moreover, studies of patients with neuropsychiatric disease that carry single-allele coding variants of DAT have provided compelling evidence for a possible causal link between genetic impairments in DAT function and the development of neuropsychiatric diseases (17-22). As a missing link in this emerging DAT-associated disease spectrum, we recently presented the first adult patient with *atypical* DTDS characterized by adult early-onset parkinsonism symptoms and ADHD as comorbid neuropsychiatric disease. This patient was compound heterozygous for two missense DAT variants, DAT-I312F and DAT-D421N. These variants cause a partial loss of dopamine uptake function *in vitro* (23). Since then, we have identified an additional patient with co-morbid parkinsonism and neuropsychiatric disease that carried the dominant-negative DAT-K619N mutation (24). Collectively, these data highlight the importance of understanding the molecular, cellular and circuitry substrates that link DAT-dependent DA dysfunction to both parkinsonism and psychiatric disease.

Here, we generate the first construct valid mouse model of atypical DTDS to elucidate behavioral and neurochemical changes that arise from DAT mutations associated with comorbid early-onset parkinsonism and ADHD. Indeed. such models of highly penetrant disease-mutations in key components of the DA system may deepen our understanding of the mechanisms through which DA exerts its multidimensional functions and uncover common neural substrates for the spectrum of clinical phenotypes shared by atypical DTDS, PD, and neuropsychiatric diseases. The mouse replicates the patient-derived heterozygous DAT-I312F/DAT-D421N genotype (23) and displays a phenotype with prominent changes in DA release and uptake dynamics together with extensive alterations in pre- and postsynaptic components of DA neurotransmission and a reduction in striatal dopaminergic fiber density. These changes are accompanied by a behavioral phenotype recapitulating core symptoms of the patient. Collectively, our work presents new insights into the neural mechanisms underlying atypical DTDS and points to potential mechanisms underlying co-morbidity between neuropsychiatric diseases and parkinsonism.

## RESULTS

### DAT-I312F/D421N^+/+^ mice display changes in DA regulation

Generation of the DAT-I312F/D421N^+/+^ compound heterozygous mice was achieved by crossbreeding of DAT-I312F^+/WT^ heterozygotes with DAT-D421N^+/WT^ heterozygotes (Supplementary Figure 1). To directly measure if the mutations have a functional impact on DA uptake *in vivo*, we first measured ^3^H-DA uptake using striatal synaptosomes. Relative to WT mice, the maximal uptake capacity of DAT-I312F/D421N^+/+^ mice was reduced by ∼75% (V_max_=26.7±9% of WT, P<0.01, Figure 1A), without significant changes in the apparent DA affinity (K_M_ WT =0.16±0.09 µM vs. K_M_ DAT-I312F/D421N^+/+^ =0.25±0.1 µM, P>0.05, Figure 1A), substantiating the disruptive properties of the disease-associated mutations previously described in vitro.

**Figure 1.**
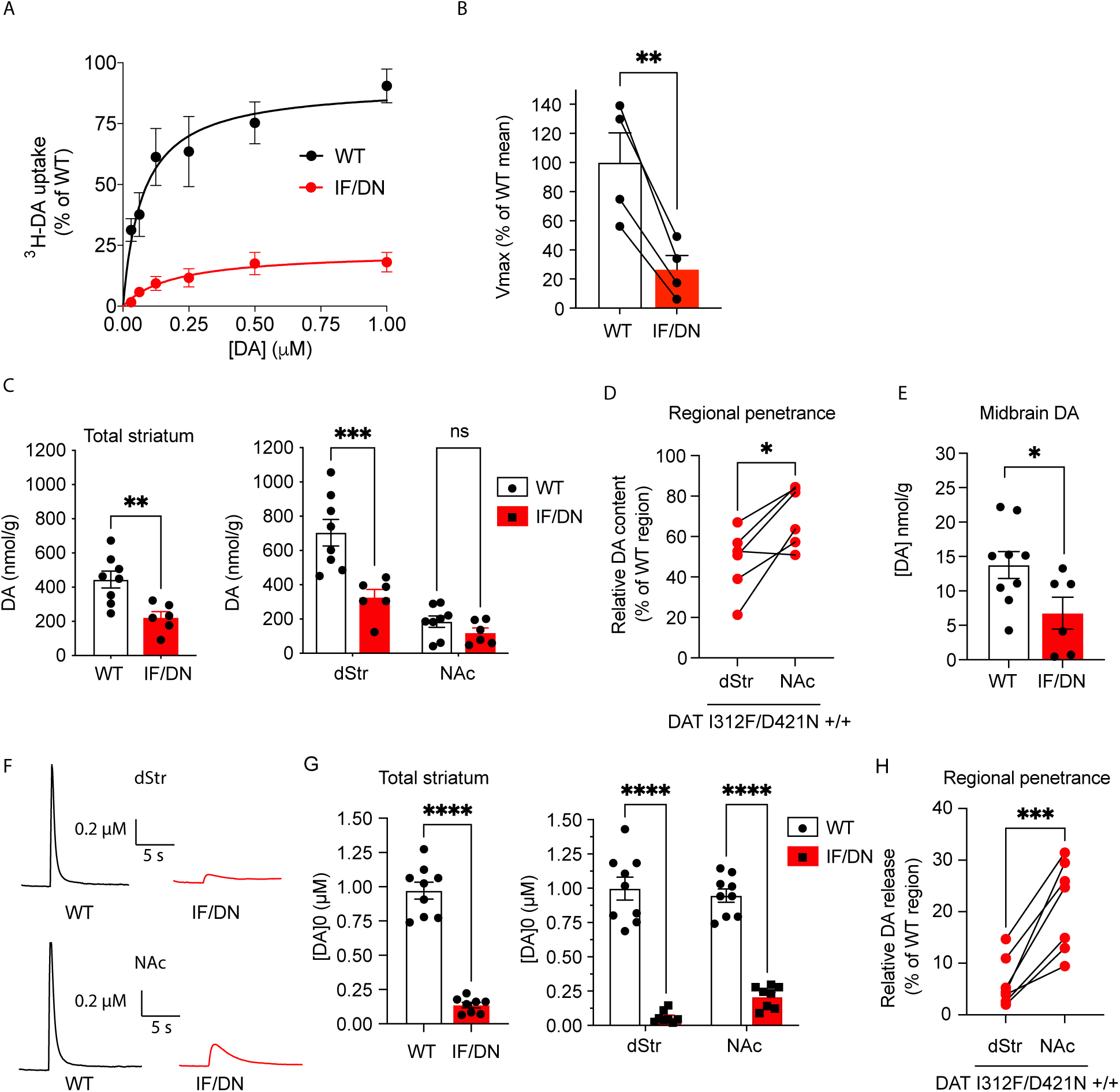
DA dysregulation in DAT-I312F/D421N^+/+^ mice. **(A, B)** ^3^H-Dopamine (DA) uptake into striatal synaptosomes from pairs of WT and DAT-I312F/D421N^+/+^ mice (IF/DN). **(A)** Average ^3^H-DA uptake curves from four experiments each normalized to the fitted V_max_ of WT mice. **(B)** Relative V_max_ of DAT-I312F/D421N^+/+^ and WT mice (**P<0.01, paired t-test, N=4 pairs of WT:IF/DN mice). There was no significant difference in K_M_ (K_M_ WT=0.16±0.09µM vs. K_M_ DAT-I312F/D421N^+/+^ =0.25±0.1µM, means±SEM, N=4, P>0.05, unpaired t-test). **(C)** HPLC analysis of striatal DA tissue content (nmol/g). Left panel, total striatal DA content (means±SEM, **P<0.01, unpaired t-test). Right panel, DA content in dStr and NAc (means±SEM, ***p<0.001 for WT_dSTR_ vs DAT-I312F/D421N^+/+^_dSTR_ and p>0.05 for WT_NAc_ vs DAT-I312F/D421N^+/+^_NAc_, 2-way ANOVA with Holm-Sidák correction for multiple testing). N=8 WT mice and 6 DAT-I312F/D421N^+/+^ mice. **(D)** Comparison of the DAT-I312F/D421N^+/+^ genotype penetrance on DA content in dStr and NAc. For each IF/DN mice, DA content in dStr and NAc was normalized to the mean of WT control mice’ corresponding regions. The impairments relative to WT are more pronounced in dStr than NAc (*p<0.05, paired t-test, N=6 DAT-I312F/D421N^+/+^ mice). **(E)** HPLC analysis of DA tissue content (nmol/g) in midbrain (means±SEM, *P<0.05, unpaired t-test, N=9 WT mice and 6 DAT-I312F/D421N^+/+^ mice). **(F)** Example traces of FSCV recorded evoked DA release from dStr (top) and NAc (bottom) in acute striatal slices from WT and I312F/D421N^+/+^ mice. **(G)** Quantification of peak DA concentrations ([DA_0_] averaged for all recording across each striatal slice (total) (means±SEM, p<0.001, unpaired t-test) and subsequently compared for the dStr and NAc regions separately (means±SEM p<0.0001, 2-way ANOVA with Holm-Sidák correction for multiple testing. **(H)** Comparison of the DAT-I312F/D421N^+/+^ genotype penetrance on DA release in dStr and NAc of DAT-I312F/D421N^+/+^ mice. Paired recordings from dStr and NAc were normalized to the corresponding means from WT mice. DA release was more impaired in dStr than NAc in DAT-I312F/D421N^+/+^ mice (p<0.001, paired t-test). Data for G-H are N=9 WT slices (6 mice) and 7 DAT-I312F/D421N^+/+^ mice slices (5 mice). *P<0.05, **P<0.01 ***P<0.001, ****P<0.0001.

Next, we measured the total DA tissue content in dorsal striatal (dStr), ventral striatal (i.e., nucleus accumbens, NAc), and midbrain samples. In general, the DA tissue content was diminished by ∼50% in DAT-I312F/D421N^+/+^ mice in both combined striatum (dStr+NAc) and midbrain (WT_STR_=440±50 nmol/g vs DAT-I312F/D421N^+/+^_STR_=220±30 nmol/g, and WT_MB_=14±2 nmol/g vs DAT-I312F/D421N^+/+^_MB_=6.8±2 nmol/g, P<0.05, Figure 1C+E). Interestingly, however, when analyzing the dStr and NAc samples separately, we found that the DA tissue content was only significantly reduced in dStr but not in NAc of DAT-I312F/D421N^+/+^ mice (WT_dStr_=700±80 nmol/g vs DAT-I312F/D421N^+/+^_dStr_=330±50 nmol/g, P<0.01, and WT_NAc_=190±30 nmol/g vs DAT-I312F/D421N^+/+^_NAc_=120±30 nmol/g, P>0.05, Figure 1C). To directly compare the regional effects of the mutations in the striatum, we normalized the DA content in dStr and NAc of DAT-I312F/D421N^+/+^ mice to the DA content in corresponding regions of WT control mice. This confirmed that dStr was significantly more affected than NAc in DAT-I312F/D421N^+/+^ mice (Figure 1D), suggesting that penetrance of the DAT-I312F/D421N^+/+^ genotype is heterogenous across striatum.

To further assess the impact of the two mutations on DA neurotransmission, we used fast-scan cyclic voltammetry (FSCV) to record stimulated DA release and reuptake in striatal slices. We carried out single-pulse stimulations with matched measurements of DA release and reuptake in dStr and NAc in each slice. Figure 1F-G show peak DA concentrations along with representative traces of the recorded DA current in WT mice and DAT-I312F/D421N^+/+^ mice. Across the striatum, evoked DA release was reduced by more than 80% in DAT-I312F/D421N^+/+^ mice compared to WT mice (WT DA_0_=0.97±0.06 µM vs DA_0=_0.14±0.02 µM in DAT-I312F/D421N^+/+^ mice, P<0.0001). Analysis of DA kinetics showed clear increases in both ‘time to peak’ (TTP, ∼2.5 fold) and halftime of clearance (T1/2, ∼5 fold) in DAT-I312F/D421N^+/+^ mice, consistent with a profound impairment in DAT function (Supplementary Table 1). Interestingly, when we compared the functional deficits from matched recordings in dStr and NAc from each DAT-I312F/D421N^+/+^ striatal slice to the corresponding regions in WT mice, DA release was significantly more compromised in dStr than in NAc (6.0±2% of WT in dStr vs 22±3% of WT in NAc, P<0.001, Figure 1F, G). Likewise, both TTP and the halftime of DA clearance were relatively longer in dStr than in NAc of DAT-I312F/D421N^+/+^ mice (Supplementary Table 1), further supporting that the impairments in DAT function have larger effects on DA dynamics in dStr than NAc. Collectively, the data show that the disease-associated DAT variants impose dramatic impairments of DA homeostasis *in vivo* with a remarkable region-specific penetrance.

### DAT-I312F/D421N^+/+^ mice show synaptopathy with terminal loss and DA receptor downregulation

Impaired DA uptake in DAT-I312F/D421N^+/+^ mice could reflect either reduced striatal protein levels or direct impairment of transport activity. To elucidate the expression and distribution of the disease-associated DAT variants, we visualized DAT in striatal and midbrain slices by immunohistochemistry. In striatal slices, we observed a clear DAT-immunosignal in both WT and DAT-I312F/D421N^+/+^ slices without a significant difference in mean labeling intensity (Figure 2A). Likewise, staining of midbrain slices showed comparable DAT immunosignal from dopaminergic neurons in both genotypes (Figure 2B). These findings suggest, in agreement with previous *in vitro* studies (23), that the DAT-I312F and DAT-D421N mutations do not cause an overall folding deficiency with ER retention and loss of striatal targeting. As a further investigation of striatal DAT protein levels, we performed western blotting on striatal synaptosomes. We observed ∼23% decrease in total DAT protein levels in DAT-I312F/D421N^+/+^ mice (P<0.05, Figure 2C), which was not apparent from the immunohistochemistry images. We also assessed the expression of tyrosine hydroxylase (TH), the rate-limiting enzyme in DA synthesis, and the vesicular monoamine transporter 2 (VMAT2) which mediates DA sequestration into synaptic vesicles. We observed a reduction in the mean TH immunosignal in striatal slices, but no change in midbrain slices from DAT-I312F/D421N^+/+^ mice compared to WT littermates (Figure 2D, E). This finding was further confirmed by western blotting of striatal synaptosomes, which showed a decrease in striatal TH expression in DAT-I312F/D421N^+/+^ mice of ∼45% (P<0.0001, Figure 2F). As observed for DAT, widefield fluorescent imaging of VMAT2 in striatal and midbrain slices showed no apparent genotype differences (Figure 2G, H), while western blotting revealed a significant reduction in striatal VMAT2 protein levels of ∼30% in I312F/D421N^+/+^ mice (73±10 % of WT, P<0.05, Figure 2I).

**Figure 2.**
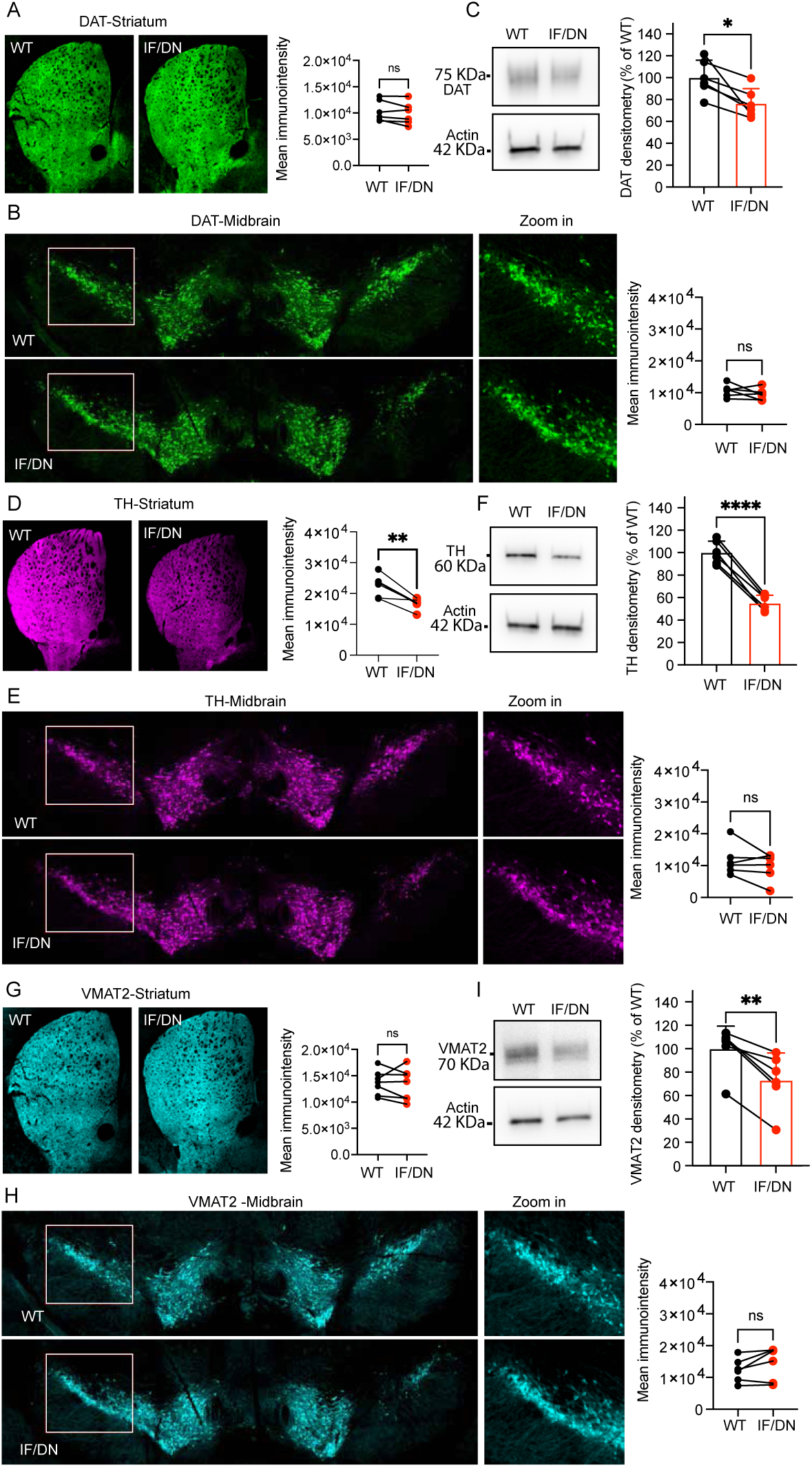
Expression of DAT, TH and VMAT2 in midbrain and striatum of DAT-I312F/D421N_+/+_ mice. Wide-field imaging of DAT in the striatum **(A)** and midbrain **(B)** by immunohistochemistry with quantification of mean fluorescent intensities shows that DAT is expressed and effectively targeted to striatal terminals in DATI312F/D421N_+/+_ (IF/DN) mice. (P>0.05, ratio-paired t-test, N=6 WT:IF/DN pairs for striatum and 5 WT:IF/DN pairs for midbrain. Every WT:IF/DN pairs were independently labelled and imaged in parallel). **(C)** Quantification of striatal DAT protein levels by western blotting of crude synaptosomal preparations shows a reduction in DAT-I312F/D421N_+/+_ mice relative to WT controls (P<0.05, ratio-paired t-test. N=6 WT:IF/DN pairs, each pair was processed and analyzed in parallel). **(D+E)** Immunohistochemical visualization of TH in the striatum **(D)** and midbrain **(E)** with quantification of mean fluorescent intensity shows a decrease in striatal TH-immunosignal in DATI312F/D421N_+/+_ mice compared to WT mice (P<0.01, N=6 WT:IF/DN pairs for striatum and P>0.05, N=5 WT:IF/DN pairs for midbrain, ratio-paired t-tests). **(F)** Quantification of striatal TH protein levels by western blotting of crude synaptosomal preparations further demonstrates that TH levels are reduced in DAT-I312F/D421N_+/+_ mice compared to WT controls (P<0.0001, ratio-paired t-test. N=6 WT:IF/DN pairs). **(G+H)** Visualization of VMAT2 with quantification of mean fluorescent intensity in the striatum **(G)** and midbrain **(H)** shows an overall comparable cellular distribution for WT and DAT-I312F/D421N_+/+_ mice (P>0.05, ratio-paired t-tests. N=6 WT:IF/DN pairs for striatum and P>0.05, N=5 WT:IF/DN pairs for midbrain). **(I)** Western blotting of VMAT2 in striatal crude synaptosomal preparations shows significantly reduced VMAT2 protein levels in DAT-I312F/D421N_+/+_ mice relative to WT controls (P<0.01, ratio-paired t-test. N=6 WT:IF/DN pairs). Data are means±SEM. *p<0.05, **P<0.01 ***P<0.001, ****P<0.0001.

The parallel reduction in striatal protein levels of DAT, VMAT2, and TH in DAT-I312F/D421N^+/+^ mice found by western blot analysis led us to investigate if the DAT-I312F/D421N^+/+^ mice display structural changes in DA terminals, which may not be resolved in widefield fluorescent images. We therefore systematically acquired confocal images from striatal slices labelled for DAT and TH to visualize the DAergic arbor in more detail. As expected, the two DAergic markers showed almost complete overlap in both WT and DAT-I312F/D421N^+/+^ mice (Figure 3A). However, the higher resolution confocal images also resolved an apparent reduction in the area density of DAergic terminals in images from DAT-I312F/D421N^+/+^ mice compared to WT mice. To directly quantify the area density of DAergic terminals, we applied a line-scan intensity analysis to count the number of terminals intersecting with a line-grid in each image (25) (see Supplementary Methods). In agreement with the visual impression, we found a significant reduction in the area density of DAT-positive terminals in the DAT-I312F/D421N^+/+^ mice compared to WT mice (P<0.001, Figure 3B). Importantly, we also found a reduction in fiber area density of TH-positive terminals (P<0.01, Figure 3C). Note that our analysis takes the lower TH expression in DAT-I312F/D421N^+/+^ mice into account as it employs a Hessian matrix that adjusts for differences in labelling intensity (Supplementary Figure 1). These data suggest that the parallel reduction in striatal levels of DAergic markers that we observed by western blot analysis may be explained, at least in part, by axonal loss in DAT-I312F/D421N^+/+^ mice and, thus, that genetic injury to DAT can compromise the microscale integrity of DAergic terminals.

**Figure 3.**
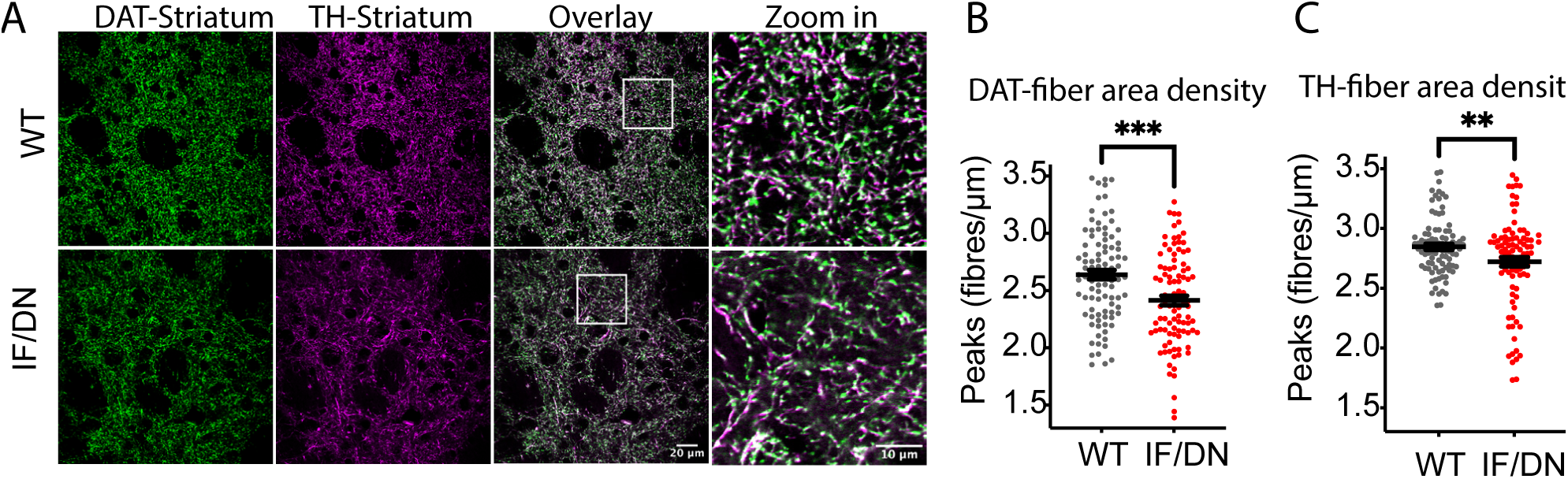
Loss of dopaminergic terminals in DAT-I312F/D421N^+/+^ mice. **(A)** Representative confocal images of striatal DA projections co-labelled for DAT and TH. **(B+C)** Quantification of area density of DAT **(B)** and TH (C)-labelled fibers. The fiber area density is reduced in DAT-I312F/D421N^+/+^ mice compared to WT (p<0.001 for DAT fiber area density and p<0.01 for TH fiber area density, unpaired t-test, N=96 WT and IF/DN images systematically sampled striatal slices from 3 independent WT:IF/DN pairs, stained and imaged in parallel, blinded to genotype. Data are means±SEM. **P<0.01 ***P<0.001.

To further investigate adaptations in the DA system, we visualized and quantified the expression of DA D1 receptors (D1R) and D2 receptors (D2R) in the striatum using knock-out validated antibodies (26, 27). Western blotting of striatal lysates showed a significant reduction in striatal protein levels of both D1R and D2R (61±8.5% and 62±12 % of WT for D1R and D2R respectively, P<0.05, Figure 4A+B). This reduction in striatal D1R and D2R levels was also observed as decreased immunolabelling intensity of striatal slices from DAT-I312F/D421N^+/+^ mice compared with WT mice (Figure 4C, D). These data strongly suggest that receptor plasticity contributes to the DAergic dysfunction in DAT-I312F/D421N^+/+^ mice.

**Figure 4.**
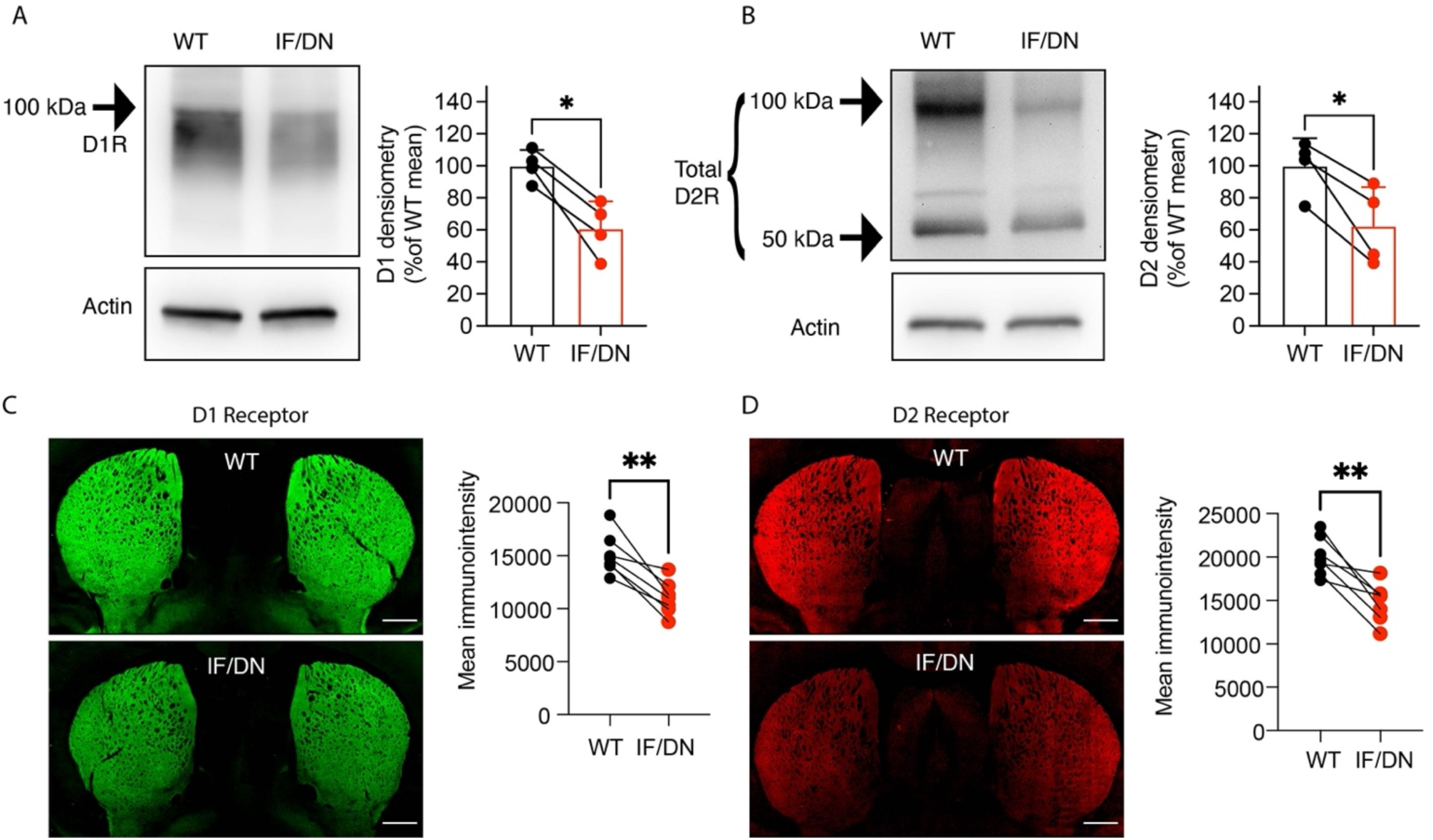
Synaptic plasticity in DAT-I312F/D421N+/+ mice. **(A+B)** Western blot analysis of DA receptor 1 (DR1) and DA receptor 2 (DR2) protein levels in striatal homogenates from pairs of WT and DAT-I312F/D421N+/+ (IF/DN) mice. The amount of both DR1 (A) and DR2 (B) is reduced in DAT-I312F/D421N+/+ mice relative to WT controls (p<0.05, ratio-paired t-test. N=4 WT:IF/DN pairs; each processed and analyzed in parallel). (C+D) Visualization of DR1 **(C)** and DR2 **(D)** by wide-field imaging of sagittal striatal brain slices from WT:IF/DN pairs (see (26, 27) for KO validation of antibodies). Visual inspection as well as quantification of mean immuno-intensity show reduced expression of DR1 (P<0.01, ratio-paired t-test) and DR2 (P<0.01, ratio-paired t-test.) in DAT-I312F/D421N+/+ mice compared to WT mice (N=6 WT:IF/DN pairs each immunolabelled and imaged in parallel). Images are representative pictures. Scale bar is 300 µm. Data are means±SEM. *p<0.05, **p<0.01.

Taken together, our findings demonstrate that the genetic insult to DAT function causes a DA dysfunction that extends beyond DA reuptake and release to also affect DA synthesis, postsynaptic receptor levels, and the microscale integrity of DAergic axonal arborization.

### DAT-I312F/D421N^+/+^ mice display persistent hyperactivity and increased explorative behavior

We next turned to characterize the overall behavioral phenotype of the DAT-I312F/D421N^+/+^ mice. These analyses were performed on adult male and female mice during a 5-week period starting at age 15±2 weeks. Of note, both male and female DAT-I312F/D421N^+/+^ mice were, by visual inspection, smaller than WT littermates. Compared to WT mice, the bodyweight was on average 4.6 grams in DAT-I312F/D421N^+/+^ males (15.6%) and 3.6 grams lower (15.2%) in females (P<0.0001, Figure 5A).

**Figure 5.**
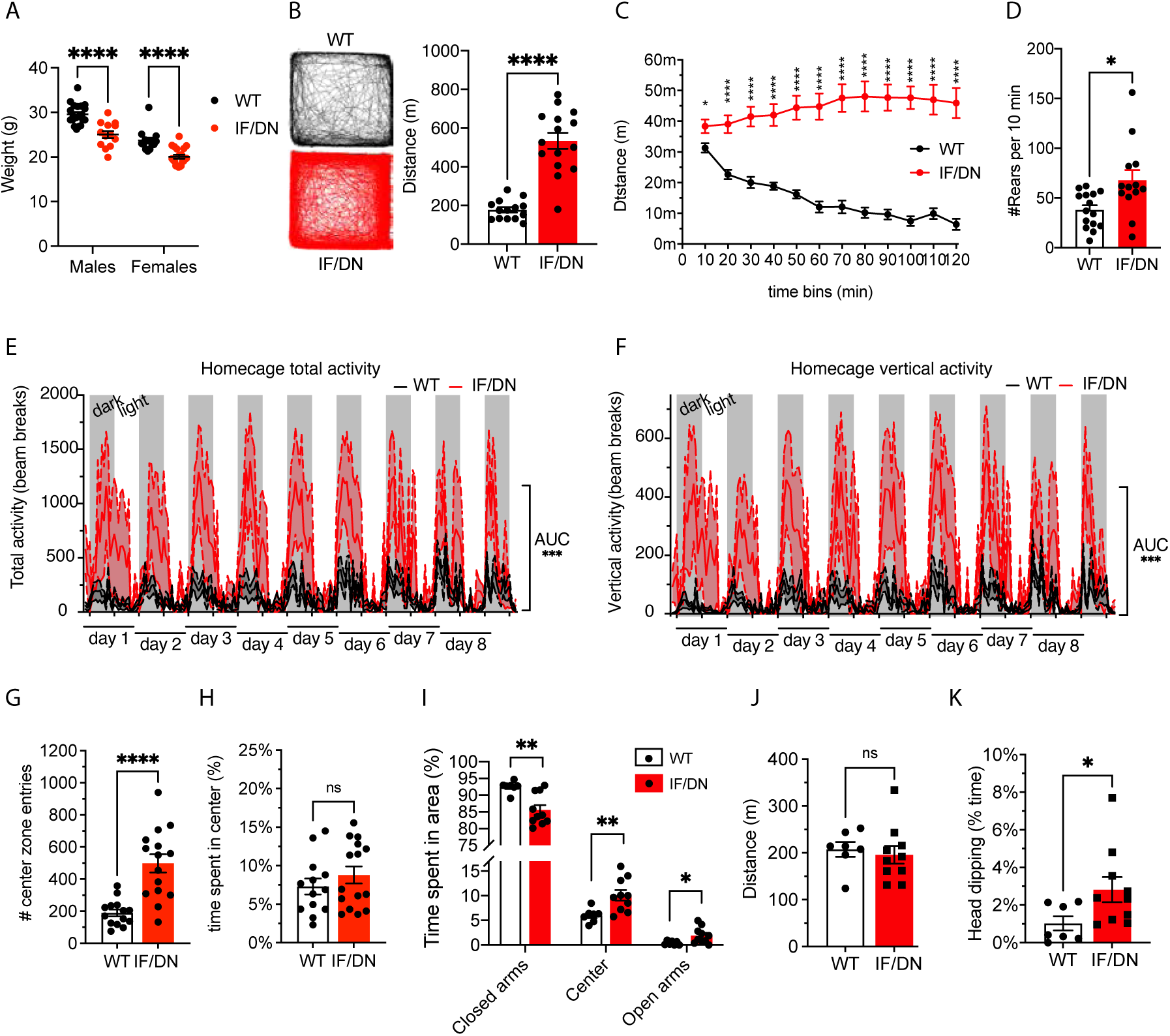
DAT-I312F/D421N^+/+^ mice display persistent hyperactivity and increased explorative behavior. (**A)** Both male and female DAT-I312F/D421N^+/+^ (IF/DN) mice have reduced bodyweight compared to WT littermates (p<0.0001, 2-way ANOVA with Holm-Sidák’s correction for multiple comparisons, N_males_= 20 WT, 13 IF/DN. N_females_= 14 WT and 20 IF/DN). (**B-C)** Locomotor activity during a 2-h open-field test. **(B)** DAT-I312F/D421N^+/+^ mice travel three times the distance of WT mice (p<0.0001, Welch’s t-test, N=15 IF/DN and 14 WT mice). **(C)** DAT-I312F/D421N^+/+^ mice show hyperactivity across all 10 min intervals during the 2-h novel open-field test. Note that WT mice habituate while the DAT-I312F/D421N^+/+^ have persistently high horizontal activity (multiple Welch’s t-test with Holm-Sidák’s correction for multiple comparisons, N=15 IF/DN and 14 WT mice). **(D)** Rearing activity during 10 min cylinder test. DAT-I312F/D421N^+/+^ mice exhibit more rears WT mice (p<0.05, Welch’s t-test, N=13 IF/DN and 15 WT mice). **(E+F)** Homecage activity recordings of total **(E)** and vertical **(F)** locomotor activity. Total activity in the homecage environment, quantified as area under the curve, is significantly higher in DAT-I312F/D421N^+/+^ mice than WT in (p<0.001, Welch’s t-test, N=6 IF/DN and 7 WT mice). **(G)** Center zone entries during 120 min of open-field recordings (p<0.0001, Welch’s t-test, N=15 IF/DN and 14 WT). **(H)** Time spend in center zone (%) during 120 min of open-field test recordings (p>0.05, Mann-Whitney test, N=15 IF/DN and 14 WT). **(I-K)** Assessment of anxiogenic versus explorative behavior in the elevated plus maze. **(I)** Time spent in closed arms, center zone, and open arms (%) demonstrate a shift towards the open arms and center zone for DAT-I312F/D421N^+/+^ mice relative to WT littermates (multiple Welch’s t-test with Holm-Sidák’s correction for multiple comparisons, N=10 IF/DN and 7 WT mice). **(J)** Distance traveled during elevated-plus maze testing (p>0.05, Welch t-test, N=10 IF/DN and 7 WT mice). **(J)** DAT-I312F/D421N^+/+^ mice spend more time dipping their heads over the sides of the open arms than WT (p<0.05, Welch’s t-test, N=10 IF/DN and 7 WT mice). All data are means±SEM. *P<0.05, **P<0.01 ***P<0.001, ****P<0.0001.

Disturbances in DAergic neurotransmission have been widely implicated in ADHD, and the patient modelled by the DAT-I312F/D421N^+/+^ mice was diagnosed with ADHD in addition to atypical parkinsonism (23). A core symptom of ADHD is hyperactivity, and we therefore compared spontaneous locomotor activity of DAT-I312F/D421N^+/+^ mice to WT littermates in a 2-h novel open-field test. On average, the DAT-I312F/D421N^+/+^ mice spent more time moving (51.4±2.6% vs 23.7±1.6%, P<0.0001) and travelled more than three times the distance of their WT littermates (534±41 m vs 177±14 m, P<0.0001 Figure 5B). This hyperlocomotive behavior of DAT-I312F/D421N^+/+^ mice was evident throughout the entire 2-h test period, but became more pronounced over time, as the WT mice habituated to the novel environment during 2 hours of recording while the DAT-I312F/D421N^+/+^ mice showed persistent high locomotor activity during the entirety of testing (Figure 5C and Supplementary movie 1-4). We compared the rearing behavior of WT and DAT-I312F/D421N^+/+^ mice in a novel environment by counting the number of rears after placing the mice in a glass cylinder for 10 min. The DAT-I312F/D421N^+/+^ mice displayed nearly twice as many rears as the WT mice (68±10 vs 38±5 rears per 10 min, P<0.05, Figure 5D), collectively demonstrating that the DAT-I312F/D421N^+/+^ mice have both increased horizontal and vertical activity in novel environments.

To assess if the observed hyperactivity was novelty-dependent, we performed extended activity recordings of DAT-I312F/D421N^+/+^ and WT mice in activity cages. During 8.5 days of monitoring, the DAT-I312F/D421N^+/+^ mice displayed persistently higher activity than WT mice both in terms of total activity (Figure 5E, P<0.001) and vertical activity (Figure 5F, P<0.001), showing that the hyperlocomotive phenotype of DAT-I312F/D421N^+/+^ mice is present even in the absence of novelty.

We next examined the effect of the DAT-I312F/D421N^+/+^ genotype on explorative versus anxiety-related phenotypes by analyzing number of center zone entries and time spend in center zone in the open field test The DAT-I312F/D421N^+/+^ mice showed a 2.6-fold increase in the number of center zone entries (498±56 entries for DAT-I312F/D421N^+/+^vs. 190±22 for WT, P<0.0001 Figure 4G), but no difference in % time spend in center zone (P>0.05 Figure 4H), indicating no anxiety-related center zone aversion. We further tested the mice for their propensity to explore versus their tendency to seek protected areas in an elevated plus maze paradigm. We found that the DAT-I312F/D421N^+/+^ mice spent more % time in the center zone and open arms than WT mice (Figure 5I). Of notice, this increase was not caused by an overall increase in locomotion, as the distance travelled by DAT-I312F/D421N^+/+^ mice during the test was not increased compared to WT mice (mean distance of 196±19 m for DAT-I312F/D421N^+/+^ vs. 208±16 m for WT mice, P>0.05 Figure 5J). Consistent with an increased explorative drive, the DAT-I312F/D421N^+/+^ mice also spent almost three-fold more time engaged in head dipping over the sides of the open arms (2.8±0.7 % of total time for DAT-I312F/D421N^+/+^ mice vs. 1.0±0.4 % for WT mice, P<0.05, Figure 5K), which is an additional behavioral correlate of explorative activity (28).

In summary, the behavioral alterations in DAT-I312F/D421N^+/+^ mice suggest that the DA imbalance imposed by the disease-associated mutations in DAT drives an ADHD-like phenotype characterized by hyperactivity and an enhanced exploratory drive.

### DAT-I312F/D421N^+/+^ mice show increased hindlimb clasping and kyphosis

Transgenic mouse models expressing genetic variants associated with hereditable forms of PD display notoriously few, if any, motor deficits (29-32). To assess the motor function of DAT-I312F/D421N^+/+^ mice, we performed a battery of motor tests comprising rotarod, pole test, hanging wire test, and hindlimb clasping during tail suspension. Most prominently, we observed a clear motor abnormality in the DAT-I312F/D421N^+/+^ mice when assessing hindlimb clasping. While DAT-WT mice showed practically no clasping behavior when lifted by their tail for 30 s, the DAT-I312F/D421N^+/+^ mice displayed clear, but varying, degrees of clasping, seen as partial or full retraction of hindlimbs towards the midline in a dystonic fashion (clasping score: DAT-WT=0.14±0.097, DAT-I312F/D421N^+/+^=9.5±1.7, Figure 6A and Supplementary movie 5-6). This clasping phenotype of DAT-I312F/D421N^+/+^ mice was not accompanied by reduced motor performance on an accelerating rotarod, in the pole test, or in the hanging wire test (Figure 6B-D), suggesting that DAT-I312F/D421N^+/+^ mice can acquire and execute motor skill to a similar level of performance as WT mice. Nonetheless, in addition to receiving higher clasping scores, the DAT-I312F/D421N^+/+^ mice also received higher scores when visually inspected for dorsal kyphosis (mean score DAT-WT=0.14±0.07, DAT-I312F/D421N^+/+^=0.59±0.1, p<0.01, Figure 6E), a phenotype which has been linked to premature aging and movement disorders (33-35).

**Figure 6.**
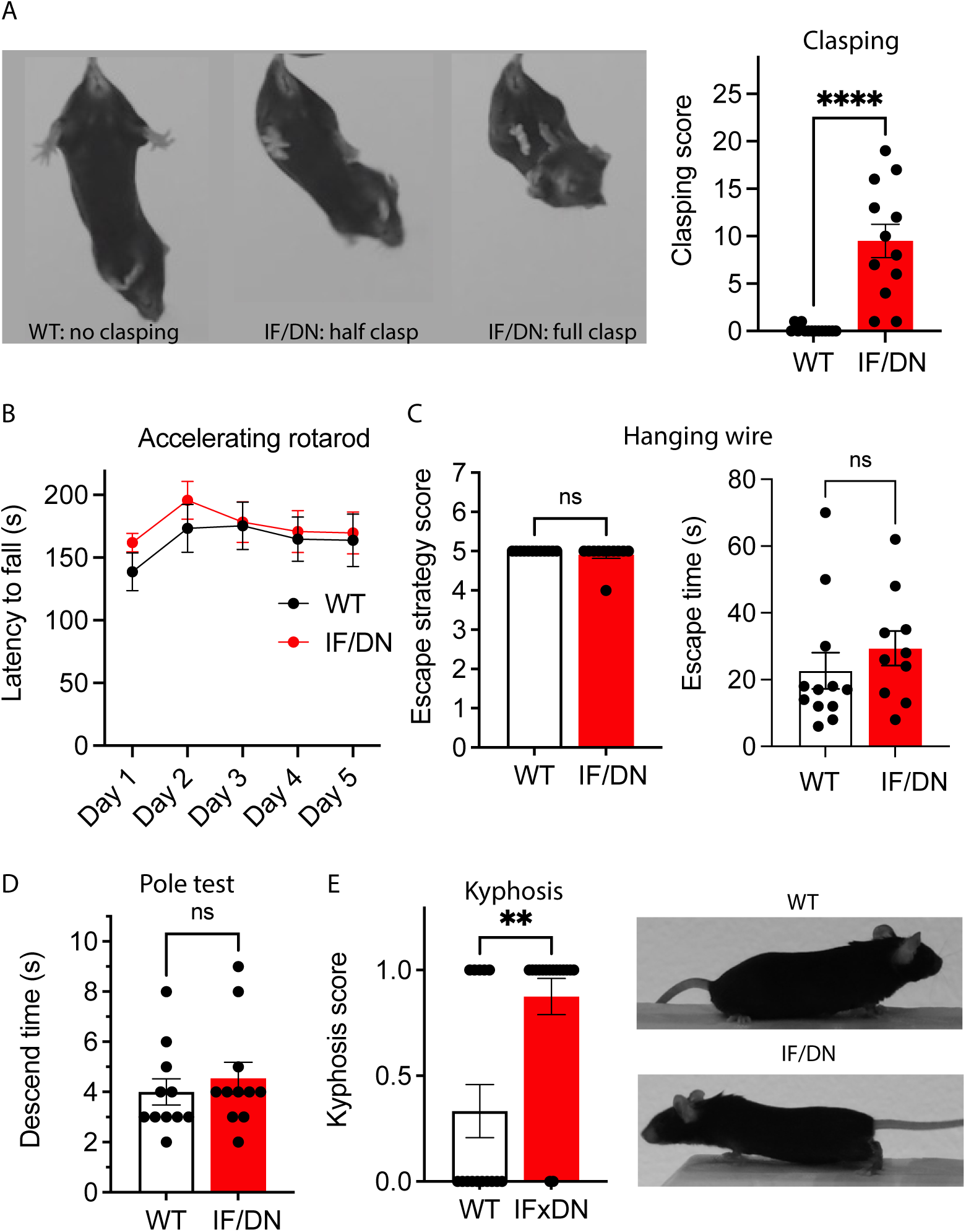
DAT-I312F/D421N+/+ mice exhibit clasping behavior and kyphosis: (**A)** DAT-I312F/D421N+/+ mice show hindlimb clasping during tail suspension. 30 sec videos were analyzed in bins of 3 sec for the presence of clasping phonotypes (see method). Compared to WT mice, DAT-I312F/D421N+/+ (KI) mice display clear clasping behavior (P<0.0001, Mann-Whitney U-test, N=14 WT and 12 IF/DN mice). **(B)** No differences were observed between WT and DAT-I312F/D421N+/+ mice in latency to fall in an accelerating rotarod test (P> 0.05 for all test days by 2-way ANOVA with Holm-Sidák multiple comparisons test). **(C)** Hanging wire test showed no difference between WT and DAT-I312F/D421N+/+ mice escape strategy scores (P>0.05, Mann-Whitney U-test, N=12 WT and 11 IF/DN mice) or in escape time to platform (P>0.05, Mann-Whitney U-test, N=12 WT and 10 IF/DN mice). **(D)** WT and DAT-I312F/D421N+/+ mice showed no difference in pole test descent time (P>0.05, Mann-Whitney U-test, N=11 WT and 11 IF/DN mice). **(E)** DAT-I312F/D421N+/+ received higher scores in visual inspection for dorsal kyphosis than WT mice (p<0.01, Mann-Whitney U-test, N=11 WT and 11 IF/DN mice). All data are mean±SEM. **P<0.01, ****P<0.0001.

### Hyperactivity in DAT-I312F/D421N^+/+^ mice is ameliorated by amphetamine and orphenadrine

Hyperactivity is a core symptom of ADHD and recapitulated in the DAT-I312F/D421N^+/+^ mouse. Since amphetamine (AMPH) is frequently used to treat ADHD in humans, we next evaluated the effect of AMPH on spontaneous locomotion in an open field arena in these mice. Following a 1-h habituation period, DAT-I312F/D421N^+/+^ mice and WT littermates were first treated with either a single dose of AMPH (2 mg/kg) or a saline injection after which activity was recorded for an additional 1-h. For AMPH-treated WT mice, we observed, as expected, a significant AMPH-induced increase in the distance traveled compared to saline controls (WT_saline_=30.9±4.1 m; WT_AMPH_=129±16 m, P<0.01, Figure 7A-B). In striking contrast, the hyperactive behavior of DAT-I312F/D421N^+/+^ mice was ameliorated by AMPH treatment, showing a reduction in travelled distance from 117±28 m in saline-treated to 34.6±7.6 m in AMPH-treated mice (p <0.01, Figure 7A-B).

**Figure 7.**
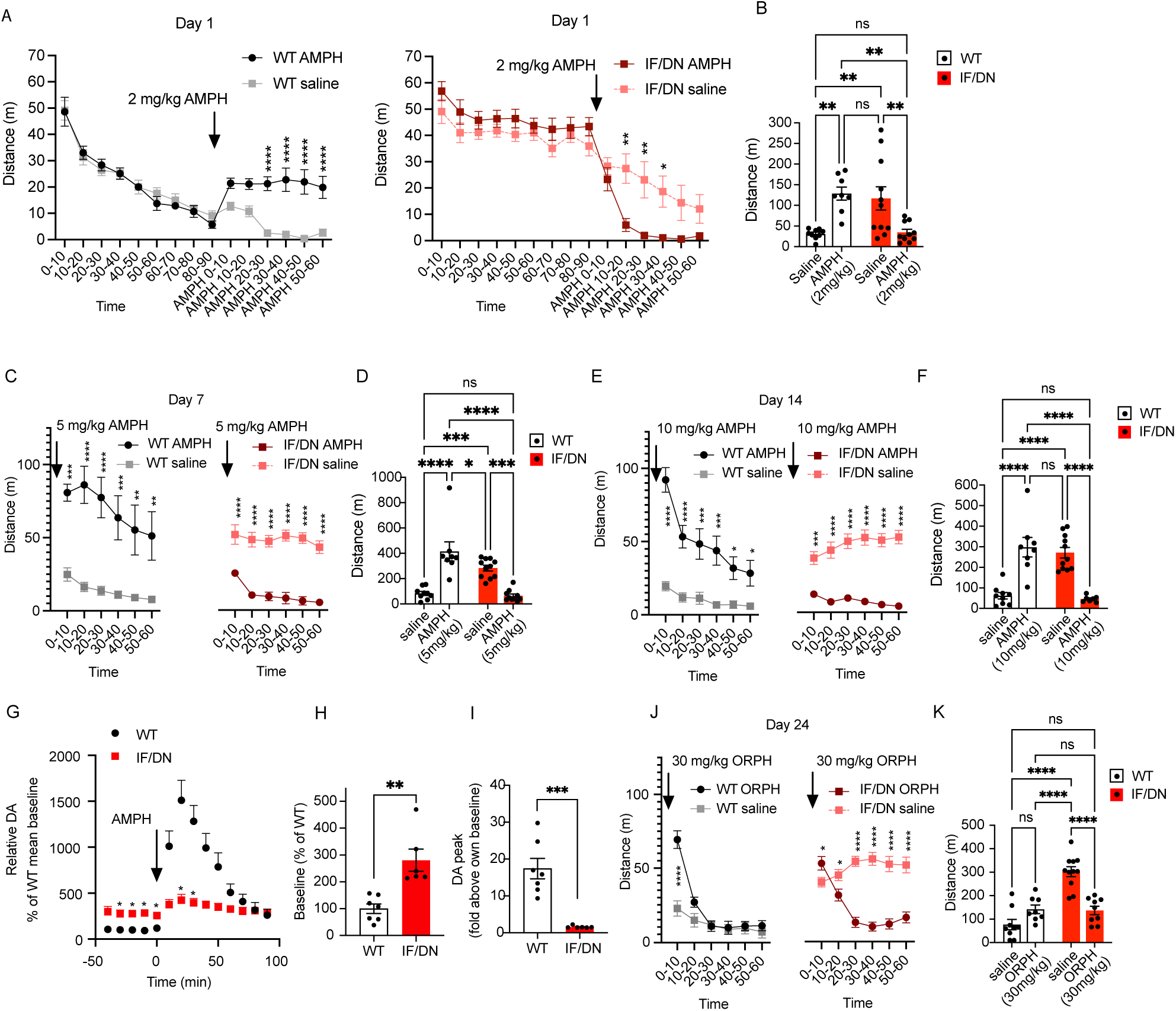
AMPH and ORPH alleviate hyperactivity in DAT-I312F/D421N+/+ mice. **(A)** Locomotor activity in an open field arena before and after administration of 2 mg/kg AMPH to WT and DAT-I312F/D421N+/+ (IF/DN) mice. Data are presented in 10 min time bins. AMPH increases locomotion in WT mice, while hyperactivity is reduced in DAT-I312F/D421N+/+ mice (repeated measure 2-way ANOVA with Holm-Sidák correction for multiple testing). **(B)** Comparison of distance travelled during 1h recording after administration of saline or AMPH (2 mg/kg) to WT and DAT-I312F/D421N+/+ mice (2-way ANOVA with Holm-Sidák correction for multiple testing, N= 9 WTsaline, 8 WTAMPH, 11 IF/DNsaline, and 10 IF/DNAMPH). **(C-F)** Re-exposure of WT and IF/DN mice to saline or 5 mg/kg AMPH (C+D) and 10 mg/kg AMPH (E+F) after 7 days washout periods. Data is shown in 10 min time bins (C+E, repeated measure 2-way ANOVA with Holm-Sidák correction for multiple testing) and as the total distance travelled during 1h recording after AMPH administration (D+F, 2-way ANOVA with Holm-Sidák correction for multiple testing, N=9 WTsaline, 11 IF/DNsaline, 8 WT5 mg/kg, 10 IF/DN5 mg/kg, 8 WT10 mg/kg, 7 IF/DN10 mg/kg). **(G)** Slow-flow microdialysis measurements of extracellular DA in the striatum of anaesthetized WT and DAT-I312F/D421N+/+ mice before and after i.p. AMPH (2 mg/kg) administration. Data is shown in % of the mean baseline for WT mice (repeated measure 2-way ANOVA with Holm-Sidák correction for multiple testing, N=7 WT and 6 IF/DN mice). **(H)** Relative baseline DA levels in % of WT mean (P<0.01, unpaired t-test, N=7 WT and 6 IF/DN mice). **(I)** Comparison AMPH-induced peak DA levels (fold above baseline) shows a larger AMPH response in WT than in IF/DN mice (P<0.001, unpaired t-test, N=7 WT and 6 IF/DN mice). **(J)** Locomotor activity in WT and DAT-I312F/D421N+/+ mice in an open field arena before and after i.p. administration of 30 mg/kg ORPH. Data are presented in 10 min time bins (repeated measure 2-way ANOVA with Holm-Sidák correction for multiple testing). **(K)** Distance travelled during 1h recording after administration of saline or ORPH (30 mg/kg) (2-way ANOVA with Holm-Sidák correction for multiple testing). N for J+K= 9 WTsaline, 8 WTORPH, 11 IF/DNsaline, 9 IF/DNORPH. All data are means±SEM. *P<0.05, **P<0.01 ***P<0.001, ****P<0.0001.

We have previously shown that both the DAT-D421N and the DAT-I312F mutations alters DAT’s substrate and inhibitor binding properties (23, 36). To exclude the possibility that higher doses of AMPH were required to elicit AMPH-induced hyperlocomotion in DAT-I312F/D421N^+/+^ mice, we re-exposed the mice to 5 mg/kg AMPH after a 1-week washout period and again to 10 mg/kg AMPH after a second 1-week washout period. Whereas WT littermates showed even more robust increases in locomotion in response to 5 mg/kg and 10 mg/kg AMPH (P<0.0001, Figure 7C-F), the DAT-I312F/D421N^+/+^ mice still demonstrated a pronounced decrease in activity at both 5 mg/kg (P<0.001) and 10 mg/kg AMPH (P<0.0001), further consolidating the attenuating effect of AMPH on the hyperactive DAT-I312F/D421N^+/+^ phenotype (Figure 7C-F).

To gain further insights into the effect of AMPH on DAT-I312F/D421N^+/+^ mice, we carried out slow-flow microdialysis experiments on anesthetized mice to compare baseline DA levels and AMPH-induced DA release in the dStr. Baseline DA levels were measured over a 40 min time window before i.p. administration of 2 mg/kg AMPH (Figure 7G). Importantly, baseline DA measurements revealed that extracellular DA levels in DAT-I312F/D421N^+/+^ mice were 2.8±0.4-fold higher than in WT mice (P<0.01, Figure 7H), demonstrating that the DAT-I312F/D421N^+/+^ genotype causes a chronic increase in extracellular DAergic tone. Administration of AMPH (2 mg/kg) elicited, as expected, a marked increase in extracellular DA in WT mice (Figure 7G+I). More surprisingly, AMPH also evoked an increase in extracellular DA levels in DAT-I312F/D421N^+/+^ mice (Figure 7G, I). However, peak DA levels increased 17±2.8 fold above baseline in WT mice whereas only a 1.6±0.13-fold increase above baseline was recorded in DAT-I312F/D421N^+/+^ mice (Figure 7I). Still, these findings suggest that the seemingly calming effect of AMPH on DAT-I312F/D421N^+/+^ mice is not established by a paradoxical decrease in extracellular DA levels leading to a reduced exploratory drive. Rather, the AMPH-treated DAT-I312F/D421N^+/+^ mice are less active despite increased extracellular DA levels.

The index patient with atypical DTDS was treated with the anticholinergic drug, orphenadrine (ORPH), to alleviate parkinsonian symptoms. We therefore decided to also compare the effect of ORPH on WT and DAT-I312F/D421N^+/+^ mice. To uncover potential disease-relevant alterations in cholinergic responses (37, 38), the effect of ORPH was evaluated in the open field test. Interestingly, analysis of locomotion in 10 min bins revealed an immediate, but short lasting (10 min) increase in locomotion in ORPH-treated (30 mg/kg) WT mice relative to saline-treated WT mice (Figure 7J). This initial hyperlocomotive response to ORPH was also seen, though less pronounced, in DAT-I312F/D421N^+/+^ mice. However, while ORPH-treated WT mice rapidly resumed activity levels similar to saline treated mice, DAT-I312F/D421N^+/+^ mice treated with ORPH showed a significant reduction in locomotor activity compared to saline control mice, an effect that persisted throughout the remaining test period (Figure 7J). Thus, during 1-h recordings, the total distance travelled by ORPH treated WT mice did not differ significantly from saline treated mice (WT_saline_=76.8±22 vs. WT_ORPH_=142±18, P>0.05, Figure 7K), while ORPH alleviated the hyperlocomotive phenotype of DAT-I312F/D421N^+/+^ mice (KI_saline_=302±21 vs. KI_ORPH_=137±18, P<0.0001, Figure 7K).

These data further support the predictive validity of the DAT-I312F/D421N^+/+^ mice, while also pointing to a potentially important cholinergic impairments that likely arises from the primary DA dysfunction.

## DISCUSSION

Parkinsonism is a multi-system neurological syndrome, characterized by bradykinesia, tremor, rigidity, and postural instability (39). The most common cause of parkinsonism is Parkinson’s disease (PD) with an estimated prevalence of 1% among people >60 years (40). The treatment of PD is symptomatic as there are no therapeutics available that can halt or delay the progressive disease course (40). Importantly, non-motor symptoms including neuropsychiatric disturbances are frequently occurring both before and co-morbid to parkinsonian features in idiopathic PD as well as in familial PD; however, our understanding of the molecular and cellular underpinnings of this comorbidity is still lacking. At present, no single animal disease model fully recapitulates the characteristics of PD, and even fewer models have been developed or evaluated as models of parkinsonism with neuropsychiatric comorbidity that show construct validity (29-32). Indeed, when considering how PD models, based on familial disease forms, continue to generate disease insights and inspire new therapeutic strategies for idiopathic and/or familial PD (41-43), the unmet need for strong models for neuropsychiatric diseases like ADHD, based on highly penetrant human mutations, is evident (44).

Atypical dopamine transporter deficiency syndrome (DTDS) is a recently described form of DTDS where the clinical phenotypes show a larger overlap with PD than classical DTDS, as it is distinct by normal motor development in infancy and early childhood. In atypical DTDS, motor deficits manifest in adolescence or adulthood and are preceded by neuropsychiatric disease in cases with adult-onset of disease (23, 24, 45). The investigation of DAT variants linked to atypical DTDS may accordingly provide key insights into how DAergic neural mechanisms are influenced by DAT and how DAergic deficits are linked to behavioral changes. In this study, we present the first mouse model of atypical DTDS based on DAT variants identified in atypical DTDS patient suffering from early-onset parkinsonism and ADHD (23). We show that the compound heterozygous genotype of the mice (DAT-I312F/D421N^+/+^) confers DAT dysfunction *in vivo*, and we explore the neurochemical disturbances that arise as a consequence and how these translate into psychomotor phenotypes that recapitulate core features of atypical DTDS. In the mice, DAT uptake capacity is reduced by ∼75%, which appears to drive a global disturbance of DA homeostasis characterized by elevated basal levels of extracellular DA, compromised DA release and spatiotemporal control, as well as an overall state of reduced DA tissue content. Immunohistochemical stainings show clear striatal DAT expression, suggesting that reduced catalytic activity underlies the loss of uptake function. This aligns with previous data from heterologous cells that also showed a marked loss of DA uptake despite efficient surface expression of the two mutants (23, 24).

Our analyses further demonstrated reduced expression of both TH and of the DA receptors (D1 and D2). It is possible that the elevated extracellular DA levels in DAT-I312F/D421N^+/+^ causes a downregulation of DA receptors, as well as reduced TH in DAergic neurons via increased tonic stimulation of presynaptic inhibitory D2 receptors. Downregulation of TH could in turn exacerbate or act as a key driver of the intracellular DA deficiency, which is already compromised by the impaired DA reuptake via DAT. Notably, our FSCV recordings revealed region-specific effects of the mutations with more pronounced impairment of release and uptake kinetics in dStr than in NAc. This is particularly interesting as it has become increasingly clear that subdomain heterogeneity within the striatum affects behavioral outcomes through differential dopaminergic signaling (46, 47). The regional differences might be a consequence of DAT being expressed at higher levels in DAergic neurons from substantia nigra, projecting predominantly to dStr, than in DAergic midbrain neurons that mainly target the NAc (48). Accordingly, dStr likely relies more heavily on DAT for maintaining intracellular DA stores and executes a tighter DAT-dependent regulation of DA transients (49).

Genetic mouse models of familial forms of PD are challenged by the general absence of clear motor impairments and DAergic neurodegeneration, unless additional genetic or exogenously strategies are employed to exacerbate the phenotype (30, 50). The ‘dying back hypothesis’ proposes that DAergic neurodegeneration is preceded by loss of axon terminals and has received increasing attention as a potential early disease state in PD (51, 52). Indeed, our visualization of the striatal axonal arbor showed that the terminal density of DAergic neurons was reduced in the DAT-I312F/D421N^+/+^ mice. Although DA toxicity most often relates to the damaging effects of intracellular DA (53, 54), the elevated extracellular DA levels that we demonstrate in the DAT-I312F/D421N^+/+^ mice could be directly implicated in this via DA toxicity (43, 55). Another intriguing possibility is that the increased extracellular DA levels in I312F/D421N^+/+^ mice drives the reduction in dopaminergic fiber density via presynaptic D2 receptors, which have been shown to act as negative regulators of the dopaminergic axonal arbor (56-58). Finally, we have previously shown that both the DAT-I312F and, particularly, the DAT-D421N mutation cause unique changes in the electrophysiological properties of DAT (23, 36). These distinct molecular phenotypes could also be directly involved in neuropathological processes that add to the detrimental effects of impaired DA uptake.

Taken together, our investigations of the DA dysfunction caused by the disease-associated DAT mutations provide a highly complex picture of a pathology involving hyperdopaminergia in a global DA deficient state with both pre-and postsynaptic plasticity. Our findings demonstrate how distinct perturbations to DAT function translate into altered dopamine-regulated behaviors and drug responses and thereby support earlier studies of genetically modified DAT mice by highlighting the critical role DAT for intracellular and extracellular dopamine homeostasis (59-62). It is interesting to note that the neurochemical profile of DAT-I312F/D421N^+/+^ mice, with 25% residual DA uptake capacity, could be expected to represent an intermediate profile of homozygous and heterozygous DAT-KO mice. However, DAT-KO heterozygous mice with 50% uptake capacity show no or only discrete behavioral phenotypes (63, 64), while the DAT-I312F/D421N^+/+^ mice display clear changes in DA-regulated behaviors that recapitulate both aspects of movement disorder and ADHD. This suggests that neural mechanisms become highly deleterious when DAT activity falls below 50% in mice. It is also possible that the distinct molecular phenotypes of the DAT-I312F and DAT-D421N mutants causes unique neurophysiological disturbances that act in concert with diminished DAT activity to exacerbate the DAergic dysfunction. Most notably, the DAT-D421N mutation is located in the second sodium binding site and introduces a sodium leak current and a propensity for the inward-facing conformation that supports constitutive DA efflux *in vitro*. The DAT-I312F mutation, on the other hand, changes the substrate-coupled anion conductance (23, 36). From a structural-functional perspective, the effect of the individual mutations on DAergic neuronal physiology would be particularly interesting to study in single-mutant knock-in mice, as these cannot be separated in the compound heterozygous mice.

The DAT-I312F/D421N^+/+^ mice exhibit an explorative phenotype, manifesting as hyperactivity, excessive rearing, and reduced preference for closed arms in the elevated plus maze test. This supports the hypothesis that the DAT-I312F and DAT-D421N mutations drive aberrant ADHD-related behaviors in consequence of the dysregulated DA neurotransmission. Conceivably, the phenotype is linked to elevated levels of extracellular DA. Previous studies have linked hyperactivity and exploration to elevated levels of extracellular DA (59, 65). Indeed, our microdialysis experiments support this link, as we found an elevation in basal DA levels in DAT-I312F/D421N^+/+^ mice. However, our data also suggest that the relationship between extracellular DA and spontaneous activity is complex as the hyperactive behaviors of DAT-I312F/D421N^+/+^ mice can be alleviated by AMPH, even though the treatment increases DA levels, albeit much less than in WT mice. Presumably, a similar enigma is relevant for patients with ADHD in whom AMPH also produces a paradoxical calming effect, even though extrasynaptic DA levels are elevated by the treatment. Interestingly, the aberrant drug-responses in DAT-I312F/D421N^+/+^ mice are not restricted to drugs that target DAT. ORPH, a drug with foremost anticholinergic effect, also alleviated the hyperactive behavior of DAT-I312F/D421N^+/+^ mice in an open field arena. This observation holds translational relevance as the patient carrying the two mutations is treated with orphenadrine. It also suggests that a neurochemical dysfunction that originates in DAergic neurons can cause more widespread circuit disturbances which in turn alters cholinergic responses (37, 38). Future work should aim at uncovering cellular and circuit deficits in other neurotransmitter systems and elucidate how such changes relate to the DA dysfunction originating from DAT.

## METHODS

### Generation, breeding, housing and genotyping of DAT-I312F/D421N^+/+^ mice

The compound heterozygous DAT-I312F/D421N^+/+^ mice were obtained by heterozygous breeding of coisogenic DAT-I312F^+/WT^ females and male DAT-D421N^+/WT^ mice which were generated by genOway S.A. (Lyon, France) as described in Supplementary Information (60). Breeding cages were kept on high-fat diet (Research Diets #D12451), while offspring received regular chow food after weening. All mice were kept on a 12-hour light/12-hour dark cycle with food and water available ad libitum. Both male and female mice were used for behavioral testing and combined for analysis of genotype differences. Primers for genotyping are listed in Supplementary Methods.

### Behavioral paradigms

Behavioral testing of DAT-I312F/D421N^+/+^ mice and WT littermates was conducted on adult mice over a period of 5 weeks starting at age 15±2 weeks. All experiments were conducted in the same order and male and female mice were tested separately. Cage-changing was prohibited 3 days prior to experiments. Experiments were conducted during the light cycle, at the same time of the day (9.30-17.30) in a quiet room with indirect lighting and mice were habituated to the test room for 45 minutes prior to experiments. Equipment was cleaned with a 70% ethanol between each animal or trial.

#### Open field test

Open field tests were used evaluate spontaneous horizontal activity. Mice were placed in white open field arenas (41×41 cm area and 50 cm height) and allowed to explore freely, while being video recorded from above. Tracking and analysis were carried out EthoVision XT (Noldus) with the center zone defined as the inner 20.5 × 20.5 cm^2^.

#### Elevated plus-maze

The elevated plus-maze (arm length 35 cm, arm width 5 cm, wall height 15 cm, 50 cm above ground) was performed to assess explorative vs anxious behaviors. Prior to testing, mice were habituated to the experimenter and the test room by individual handling of each mouse for 5 min for 3 consecutive days. On day 4, mice were placed on the open arm facing the center zone and video recorded from above for 15 min. Tracking and analysis were carried out in EthoVision XT to derive: distance travelled, time spent in the open/closed arms (center point and tail point) and the center zone (center point). In addition, time spent nose-dipping from within the center zone or on the open arms were derived as an additional parameter of exploratory behavior.

#### Rearing

Mice were placed in an empty glass beaker (diam. 20cm, height 22cm) in front of two vertical mirrors arranged at an angle of approximately 110° and video recorded for 10min. Frequency of rears, defined as lifting the front paws from the ground and upwards stretching the body were manually counted.

#### Activity measures in the home cage environment

For home cage activity recordings, mice were singly-housed in metabolic cages (Phenomaster, TSE systems, Bad Homburg, Germany) with food and water ad libitum. Nesting material could not be provided as infrared sensors were used to detect movement. Horizontal locomotor activity, grooming/fine activity, and rearing were monitored by infrared beam breaks. Measurements were collected for 8.5 days without human intervention except for required visual inspections.

#### Hanging Wire Test, Pole Test, and Rotarod

To test muscle strength, agility, and motor learning and coordination, mice were subjected to the hanging wire, pole test, and rotarod as described in Supplementary Methods.

#### Clasping and kyphosis

Clasping behavior was assessed by lifting each mouse by its tail during video recording. Videos of 30s were analyzed manually at low speed in 3s time-bins (i.e., 10 time-bins per video) for clasping behaviors. For each time-bin, a score of 0 was given if no clasping behavior was seen, 1 if the mice performed a ‘half clasp’ (one hind paw fully retracted to body midline or both hind paws retracted halfway to the body midline) and 3 if a clasp was fully established (both hind paws retracted, see Figure 5). The scores for each of the 10 time-bins were combined into a clasping score (hence the max score is 30).

To assess presence of abnormal anteflexion of the spine (kyphosis), mice were placed on a straight surface and observed with video recording during exploration for 60s. Kyphosis was scored as 0=not present, 1=mildly present and can be stretched out, 2=obviously present and cannot be stretched out.

### Microdialysis

Mice were pretreated with the analgesic carprofen (5 mg/kg in 10 ml/kg s.c.; Pfizer, New York, NY), anesthetized with isoflurane (induction ∼4%, maintenance ∼1%; Baxter, Deerfield, IL) and placed in a stereotaxic frame (Kopf Instruments, Tujunga, CA). A microdialysis probe (CMA7, 2 mm, CMA microdialysis, Kista, Sweden) was inserted into the dStr at bregma coordinates: AP: 1.2 mm, ML: ±1.5 mm and DV -4 mm (tip position relative to skull). The probe was connected to a syringe pump and perfused with artificial cerebrospinal fluid (aCSF; Harvard apparatus, Holliston, MA). Microdialysis was performed in anesthetized mice with a flow rate of 1.2 μl/min. Microdialysates were collected every 10 min and analyzed immediately for DA content on a HTEC-500 HPLC (Eicom, Kyoto, Japan), separated on a PP-ODS affinity column (Eicom) and electrochemically detected. Chromatograms were analyzed using the EPC-300 software. After 40 min of stable baseline measurements, AMPH (2 mg/kg) was injected i.p and microdialysates were collected for another 90 min. DA standards (0.5 pg/μl and 10 pg/μl) were run before and after microdialysate measurements. Relative DA levels (AUC normalized to WT mean baseline) were used to depict the time course of AMPH-induced DA release and for comparing average baseline levels in WT and DAT-I312F/D421N^+/+^ mice. For each mouse, the baseline DA level was determined as the mean of the four samples proceeding AMPH administration. AMPH-induced DA efflux was quantified by the DA peak levels derived as the ‘fold increase’ above own baseline.

### Synaptosomal uptake

DA uptake on synaptosomal preparations were carried out as described in (66). All experiments were performed on pairs of WT and DAT-I312F/D421N^+/+^ mice. All pairs were independently processed and matched WT and DAT-I312F/D421N^+/+^ samples were run in parallel during the entire experiment. A two-fold dilution row (final concentrations 1-0.031 µM) containing a mixture of unlabeled DA and 2, 5, 6-[^3^H]-DA (Perkin Elmer Life Sciences, USA) was used to obtain saturation DA uptake curves, which were fitted with Michaelis-Menten kinetics to derive V_max_ and K_M_ values.

### Immunohistochemistry

Immunocytochemistry was performed on coronal sections of the striatum and midbrain from transcardially perfused mice (4% paraformaldehyde in 0.1 M PBS (pH 7.4)) as described in (24). WT and DAT-I312F/D421N^+/+^ derived slices were stained in pairs and imaged in parallel. Primary antibodies (see Supplementary Table 2) against TH, DAT, VMAT2, DR1 (see (26) for KO validation) and DR2 (see (27) for KO validation) were incubated overnight. Secondary Alexa488, Alexa568, and Alexa647-conjugated antibodies (1:400, Abcam) were incubated either ON 4°C or 3h at RT. Sections were mounted on glass coverslips (Mentzel-Gläzer, 24×60 mm, Germany) using ProLong Gold antifade mounting media (Invitrogen)

### Fluorescent imaging and image analysis

Wide-field fluorescent images of entire striatal and midbrain sections were acquired using a Slide Scanner Axio Scan.Z1 with a Plan-Apochromat 20x/0.8 objective (Zeiss). Alexa-488, Alexa-568, and Alexa-647 emissions were recorded using a 525/50 nm band pass (Alexa 488), 605/70 nm band pass (Alexa 568) and 690/50 nm band pass (Alexa 647) filters, with an identical exposure setting for WT and DAT-I312F/D421N^+/+^ mice. Image processing and analysis of immunoreactivity was carried out using Fiji software (NIH ImageJ2). Nonspecific signal was subtracted from each image and rectangular ROIs around the entire midbrain or bilateral striatae were defined. Automated threshold functions were applied for unbiased isolation of labelled structures (‘Li threshold’ function for striatal images, ‘mean threshold’ for TH midbrain and ‘default threshold’ function for DAT and VMAT2 midbrain stains). The mean signal intensity of the threshold area was measured within each ROI and used to compare the signal intensities in WT and DAT-I312F/D421N^+/+^ sections.

Confocal images of DAT and TH were used for analysis of DAergic terminal area density using a line-scan intensity analysis, modified from (25). This analysis accounts for potential differences in protein expression as described in detail in the Supplementary Methods. Confocal images were acquired on a LSM 510 confocal laser-scanning microscope with an oil immersion 63x/1.4 numerical aperture objective (Carl Zeiss, Oberkochen, Germany), using identical settings for matched WT:IF/DN pairs. Imaging of WT:IF/DN pairs stained in parallel was done on the same day, blinded to the genotype. Images with regional comparability were systematically collected throughout each striatal slice. A total of 32 none-overlapping images were acquired from each mouse. Alexa-488 dye was excited using an argon-krypton, laser and the emitted light was detected using a 505-530-nm band pass. A 543-nm helium neon laser was used to excite the Alexa-568 fluorophores, and fluorescence was obtained using a 560-nm long-pass filter.

### Western blotting

Striatal tissue lysates and crude striatal synaptosomes were made as previously described (24, 67). After adjustments of protein concentrations, equal amounts of protein were resolved by SDS-PAGE and Western blotting was carried out as described in (23, 24) using primary antibodies (against TH, DAT, VMAT2, DR1 and DR2 (see Supplementary Table 2). Blots were developed by chemiluminescence following incubation with secondary HRP-conjugated antibodies (1:2000) and an HRP-conjugated anti-β-actin antibody (1:40.000, Sigma) was used as loading control. Relative band intensities were quantified using Fiji software.

### Fast-scan cyclic voltammetry

Fast scan cyclic voltammetry recordings were made on acute brain slices from adult DAT-I312F/D421N^+/+^ mice and WT littermates. The animals were anesthetized with halothane, quickly decapitated and the brain harvested. Next, the brain was submersed in ice-cold oxygenated artificial cerebrospinal fluid (aCSF) containing (in mM): NaCl (125), KCl (2.5), KH2PO4 (0.3), NaHCO3 (26), glucose (10), CaCl2 (2.4), MgSO4 (1.3) and coronal striatal brain slices of 300 µm thickness were prepared with a VT1000S vibrating blade microtome. Slices were transferred to oxygenated aCSF at room temperature and allowed to recover for at least 1h. Single-pulse (1 ms, 400 µA) evoked release of DA in dStr and NAc was recorded by fast-scan cyclic voltammetry (FSCV) as previously described (57).

### HPLC analysis of total DA tissue content

Mice brains were dissected to obtain dStr, NAc, and midbrain samples, which were snap-frozen in liquid nitrogen immediately after dissection and kept at -80°C until processed. The tissue samples were homogenized using 300 µL of lysis buffer (20 mM HEPES, 125 mM NaCl, 10% Glycerol, 1 mM EDTA, 1mM EGTA) supplemented with 1% protease inhibitor cocktail (Millipore), and 30 µL 1N HClO4. The homogenate was centrifuged for 15 min at 16,000 g (4°C) and the supernatant was collected and filtered through a 0.22 µm filter. DA content was analyzed using high-performance liquid chromatography with electrochemical detection (HPLC-ECD) as previously described (68). DA content was normalized to the total protein in each sample.

### Statistics

GraphPad Prism 8.0 (GraphPad Software, San Diego, CA) software was used for statistical analysis and data fitting. All, statistical methods are specified in the figure legends. Data are presented as mean±SEM. Paired or unpaired (depending on the experimental design) two-tailed t-test were applied for 2-group comparisons for normally distributed data. Mann-Whitney (unpaired) or Wilcoxon (paired) t-tests were used for data that failed normality tests. 2-way ANOVA with Holm-Sidák posttest were used for multiple comparisons. Differences were considered significant for p<0.05.

## Study approval

All animal experiments were approved by the Danish Animal Experimentation Inspectorate (permission numbers 2017-15-0201-01160 and 2017-15-0201-01177) and adhere to the European guidelines for the care and use of laboratory animals, EU directive 2010/63/EU.

## Author contributions

F.H. and U.G. conceptualized the study. F.H., L.K.K., C.P., B.D-L. C.F.R., N.A., J.P.R., and F.B. conducted the experiments. F.H., U.G., M.R., G.T., L-E.T., B.H., and D.W., provided supervision. F.H., and U.G. prepared the final figures and wrote the original manuscript. F.H. and U.G. provided funding. All authors contributed to the editing and review of the manuscript.

## Acknowledgement

We thank Anette Dencker Kaas for excellent technical assistance. The work was supported by: Independent Research Fund Denmark – Medical Sciences (DFF-4183-00571, FH; DFF 4004-00097B, UG), Lundbeck Foundation (R181-2014-3090 and R303-2018-3540, FH), Lundbeck Foundation (R223-2016-261, UG), Fondecyt Initiation Fund (11191049, JAP), Canadian Institutes of Health Research (PJT-165928 and PJT-183931, LT), the Krembil Foundation (LT), and the Aligning Science Across Parkinson’s (ASAP) initiative (LT).

