## Supplementary information for "Mouse model of atypical DAT deficiency syndrome uncovers dopamine dysfunction associated with parkinsonism and psychiatric disease" (from another version of this preprint: version 1, 2023-08-17)

---

### SUPPLEMENTARY MATERIALS AND METHODS

#### Generation of DAT-I312F/D421N$^{+/+}$ mice

The compound heterozygous DAT-I312F/D421N$^{+/+}$ mice were generated by heterozygous breeding of DAT-I312F and DAT-D421N mice strains (Supplementary Figure 1), which were made by genOway S.A. (Lyon, France). Homology recombination was used to introduce the hDAT-I312F (mDAT-I311F) and the hDAT-D421N mutation (mDAT-D420N) in the murine *Slc6a3* gene for expression under the control of the endogenous *Slc6a3* promoter. The targeting vector for the DAT I311F mutant line contained a murine *Slc6a3* fragment spanning from intron 4 to intron 8 with the point mutation I311F (ATC to TTC) in exon 7 and a neomycin cassette flanked by loxP sites in intron 6. The targeting vector for generation of the DAT-D421N mouse line contained a murine *Slc6a3* region from intron 8 to intron 13, including the point mutation D420N (GAC to AAC) in exon 9 and a loxP-flanked Neomycin cassette in intron 9.

Following linearization, each targeting vector was electroporated into ES cells derived from black-coated C57BL/6J mice and subjected to positive selection. PCR sequencing and southern blotting was used to identify ES cell clones haboring correct recombination events. Recombined ES cell clones, (from black-coated C57BL/6 mice), were injected into blastocysts, derived from an albino C57BL/6 mice strain and implanted in uteri of pseudopregnants females to produce chimeras. Chimeras were identified by coat color and male chimeras (>50%) were bred with Cre-recombinase expressing C57BL/6 mice to excise the *lox*P flanked neomycin cassette. Resulting heterozygous mice carrying the neo-excised point mutant knock-in allele were validated by PCR genotyping and by Southern blotting to confirm successful recombinase-mediated excision of the neomycin cassette.

#### Genotyping of DAT-I312F/D421N$^{+/+}$ mice

Ear or tail biopsies were used for PCR genotyping using the following primers for detection of the I312F mutation: 5' - CTAGGGATGCGGTAGGGCTTGTTTC - 3' and 5' - CTGGGGATGCTTCATTTCCAAGAGG - 3', and for detection of the D421N mutation: 5' - GGACATGAGTGTTAGTGGCTGGAATGTAAC - 3' and 5' - CTCAGTAAATAATCTTGAGCAGTTGATGGGG - 3'.

**Hanging Wire Test**

To test muscle strength and coordination, mice were subjected to the hanging wire test (1). Mice were placed, hanging by their front paws in the middle of an affixed horizontal metal wire (2mm thick and 40cm long) 37cm above a cage filled with cushioning bedding with escape platforms on both ends of the wire. All mice were video recorded during a maximum of 3 trials (intertrial period 10 min) with a maximum trial length of 300 seconds. The performance was scored manually as follows: 0 = fell off; 1 = hung onto the bar with two forepaws; 2 = hung onto the bar with added attempt to climb onto the bar; 3 = hung onto the bar with two forepaws and one or both hind paws; 4 = hung onto the bar with all four paws and with tail wrapped around the bar; and 5 = grips the bar with all four paws and with tail wrapped around the bar and escapes to the platform; -1 = improper behavior. Improper behavior, such as balancing on or deliberately letting go of the wire, was addressed by immediately placing the mouse back on the wire. In addition to the scoring of escape strategy we also analyzed 'time on the wire'. If the mouse fell from the wire the experiment was repeated for a maximum of 3 trials and the best score was used. The best score was used for statistical analysis.

**Pole Test**

To assess agility, coordination and ability to descend headfirst, we performed the pole test. Animals were trained for 5 trials on two consecutive days and tested with video recording for 5 trials on the third day. A wooden pole wrapped in gauze (50cm, 1cm diameter) was placed in the center of the home-cage with the base covered in bedding material. Mice were placed at the top of the pole with the head facing upwards and time to descend was measured. Median data across 5 trials from video recordings was used for the analysis.

**Rotarod**

An accelerating rotarod (Panlab/Harvard Apparatus, USA; rod diameter 3.5cm) was used to assess motor learning and coordination. Mice were first trained to stay on the rotating for 180s at 5 rpm. Mice were then tested with an accelerating protocol (4-40rpm in 5min) for 5 consecutive days with 3 trials per day (10 min inter-trial period). The time to fall was recorded and time was also stopped if a mouse completed two passive rotations. The best performance on each day was used for statistical analysis.

**Striatal fiber area density analysis**

To quantify fiber area density from confocal images of dopaminergic projections in striatum of WT and DAT-I312F/D421N$^{+/+}$ mice, we implemented a line-scan intensity analysis inspired by (2). To minimise the risk that potential differences in terminal density reflected differences protein expression, rather than true terminal loss, all images were first fitted with a Hessian-based image filter in Image J (version 2.0.0, plugin, Feature J (Meijering, 2003)). This filter uses the Hessian matrix to find local intensity variations – i.e. maxima (fluorescent fibers) and minima (background) - and enhance their extremities to extract the curvilinear structures of the labelled fibers. The images were then converted to a binary image for subsequent analysis of fiber density. When converting images, "smoothing scale factor" was set to 0.5 and the "absolute eigenvalue comparison" option was chosen, generating a resulting image of largest eigenvalues (extrema) – settings were chosen in conjunction with (2). A line-scan intensity analysis was then carried out on each image using a grid of 9 vertical and 9 horizontal lines of same length (202.031μm) and equidistant apart (lines created conformed to a set grid function in ImageJ) for an unbiased and automated analysis of the number of fibers crossing each line (Supplementary Figure 2). The pixel intensity along each line was plotted using ImageJ's "Plot Profile" tool (Supplementary Figure 2D). Only lines fully contained within the striatal region were profiled.

Next the line intensity profile data were imported to MATLAB (version 2018b 9.5.0.944444; MathWorks). Line scans were baseline-adjusted using the "msbackadj" command from MATLAB's bioinformatics toolbox. This algorithm determines the baseline by classifying data points into peaks and background, with the mean value of the background class used as the estimated baseline for each line scan. The resulting intensity matrices with adjusted baselines were then processed through a peak detection algorithm; "mspeaks" (Supplementary Figure 3). More detailed information on both processing commands are available in (3). The average background intensity from all line scans over the entire striatum between stained WT/KI pairs were used as a height/threshold filter. Local maxima above the threshold were counted as peaks and used to represent an individual fibres crossing the line scan. The average number of peaks per line scan in each image was divided by the length of each line (202.031μm) to yield the mean value of peaks/fibers per μm. The results for WT and DAT-I312F/D421N$^{+/+}$ mice were then compared.

**SUPPLEMENTARY FIGURES:**

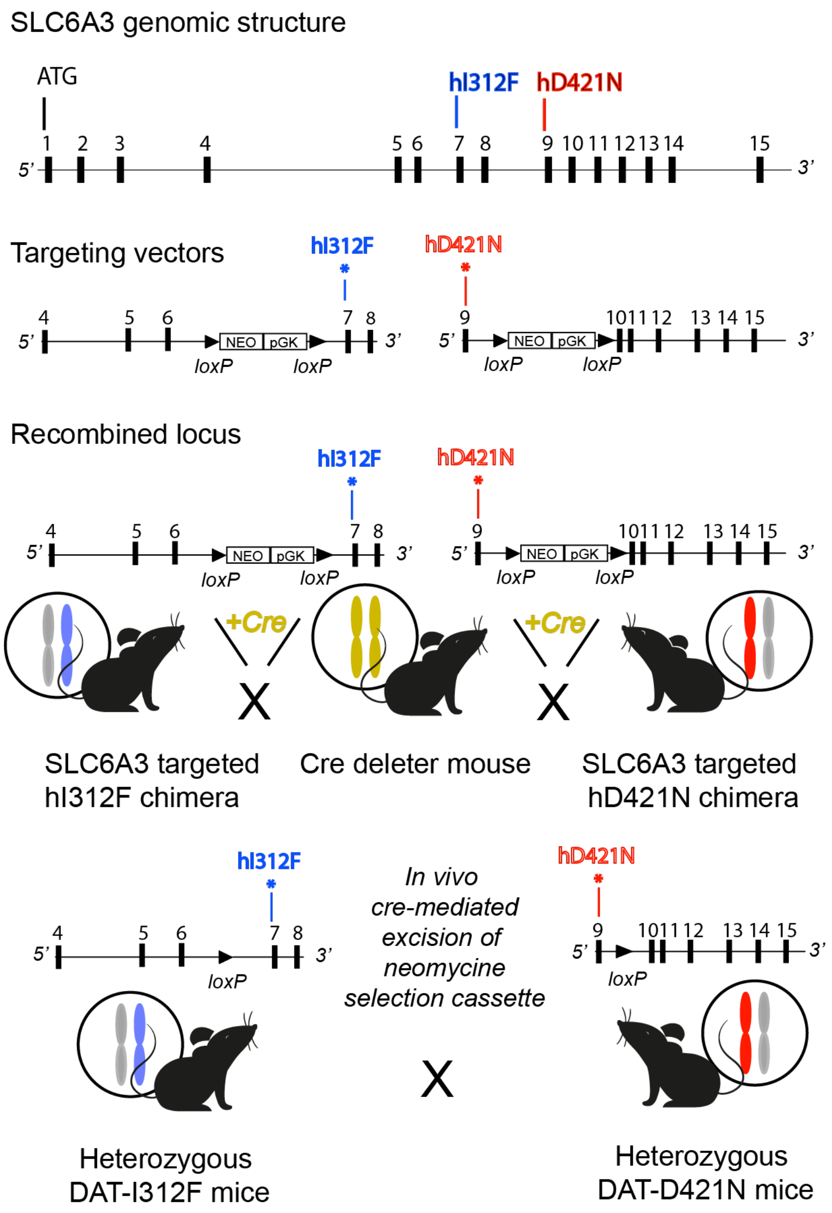


**Supplementary Figure 1. Generation of transgenic DAT-I312F/D421N$^{+/+}$ mice.** Homology recombination was used to introduce the hDAT-I312F (mDAT-I311F) and the hDAT-D421N mutation (mDAT-D420N) in exon 7 and 9 of the murine *Slc6a3* gene for expression under the control of the endogenous *Slc6a3* promoter. The DAT I311F targeting vector contained a murine *Slc6a3* fragment spanning from intron 4 to intron 8 with the point mutation I311F in exon 7 and a neomycin cassette flanked by loxP sites in intron 6. The targeting vector for generation of the DAT-D421N mouse line contained a murine *Slc6a3* region from intron 8 to intron 13 with the D420N point mutation in exon 9 and a loxP-flanked Neomycin cassette in intron 9. Male chimeric mice were bred with Cre-recombinase expressing C57BL/6 mice to excise the *lox*P flanked neomycin cassette. Female DAT-I312F and male DAT-D421N heterozygous mice carrying the neo-excised point mutant knock-in allele were cross-bread to generate compound heterozygous DAT-I312F/D421N$^{+/+}$ mice.

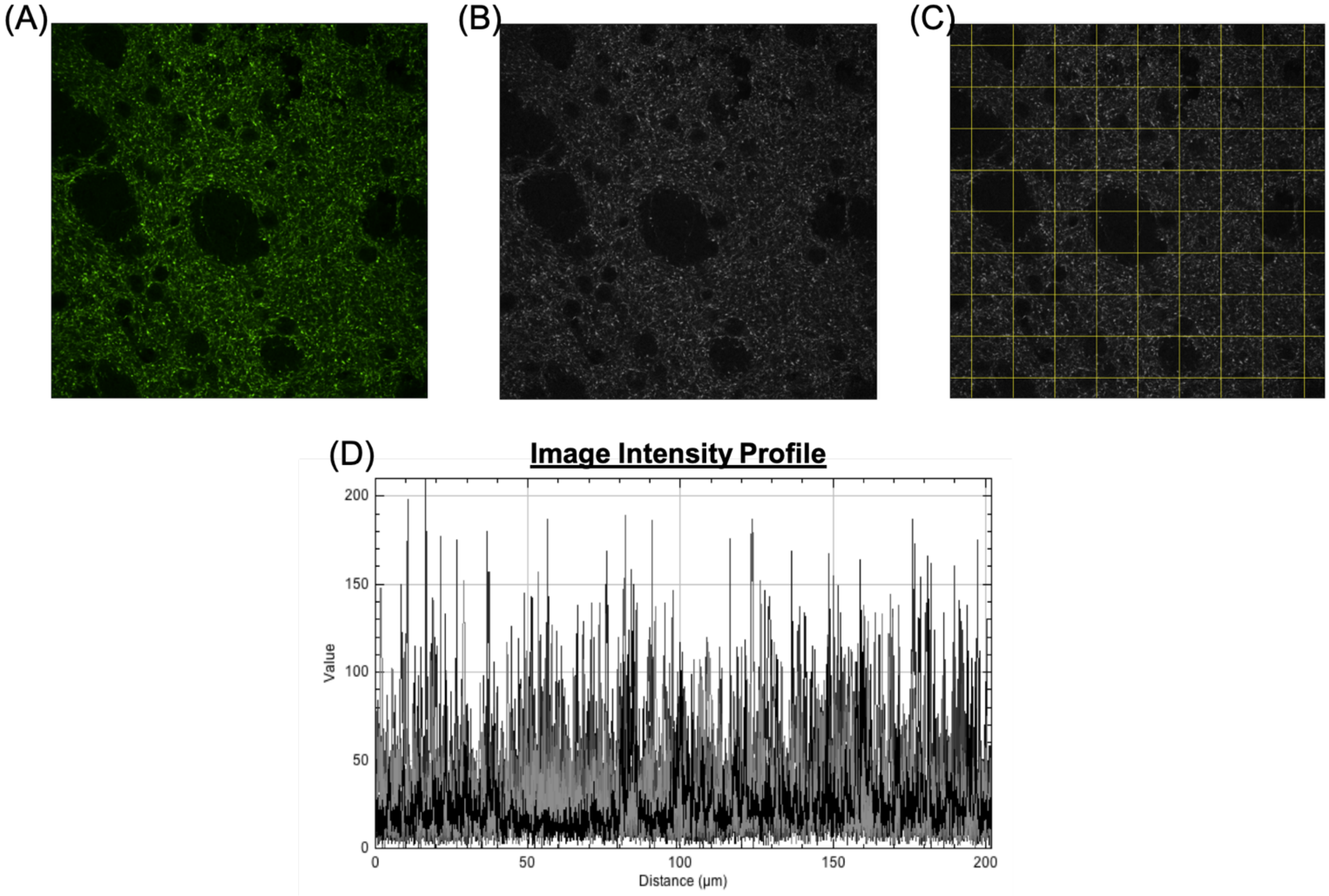


**Supplementary Figure 2. Example for the application of Hessian-based edge detection and image intensity profiling in ImageJ. (**A). Example image from of DAT-labelled dopamine fibers in striatum (B). Application of Hessian-based filter on image in order to extract a second order curvilinear structure from the image. (C). Line ROIs drawn, conforming to set grid function in ImageJ. All lines remain in the striatal region. (D). ImageJ's "Plot Profile" tool plots the pixel intensities along the length of each line (202.031 μm in length). The given graph displays each line's intensity profile stacked on top of each other. The XY coordinates of the profile can inputted to MATLAB software for further analysis.

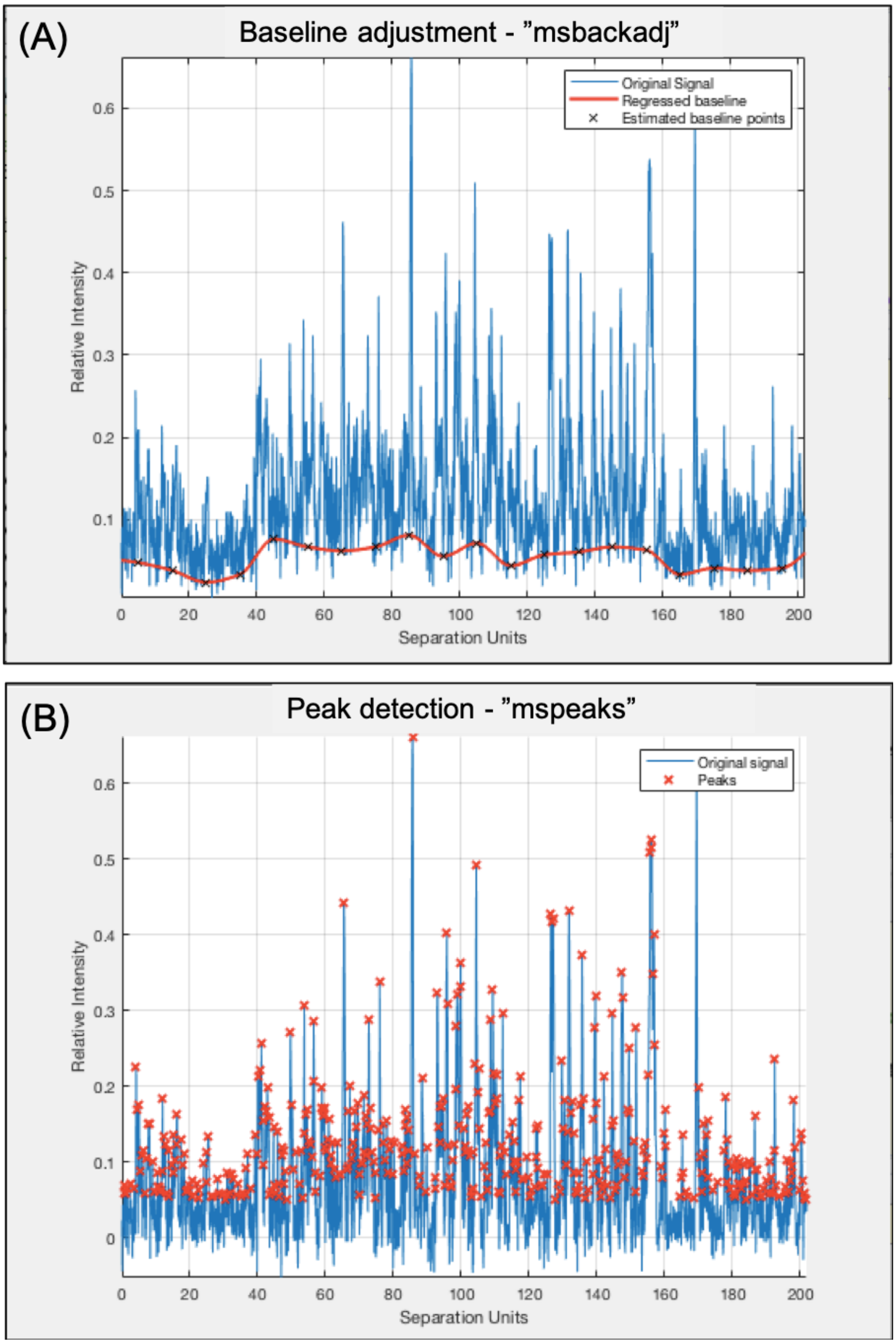


**Supplementary Figure 3. Single line scan representative examples for the functions “msbackadj” and “mspeaks”.** (A). Represented single line scan with baseline displayed after processing data using “msbackadj” function. Intensities have been normalised to the highest intensity detected in the image profiled. The “msbackadj” algorithm classifies the data into signal and background, calculating the average background/threshold in bin sizes of 10 µm (separation units). (B). The same represented line scan after baseline adjustment. Detected nerve fibres are marked as Xs (peaks) identified by the “mspeaks” algorithm. The mean of all calculated background values between each WT/KI pair acts as the calculated threshold for the image. All peaks below this threshold, such as those around unit 160 (µm), are not included in the peak detection algorithm. Local maxima are calculated as peaks, representing fibre counts.

**SUPPLEMENTARY TABLES**

**Supplementary Table 1. Kinetic parameters for DA release and reuptake in striatal slices.** Kinetic parameters (time to peak (TTP),T1/2, and DA peak) of DA neurotransmission derived from FSCV recordings of single-pulse evoked DA released in dStr and NAc from acute striatal slices. Significant impairments are found across all parameters for the DAT-I312F/D421N$^{+/+}$ mice relative to WT mice (multiple unpaired t-test with Holm-Sidák correction for multiple testing, *p<0.05, **P<0.01 ***P<0.001, ****P<0.0001.). Comparison of the DAT-I312F/D421N$^{+/+}$ genotype penetrance in dStr and NAc of DAT-I312F/D421N$^{+/+}$ mice was done by normalizing paired recordings from dStr and NAc to the corresponding means from WT mice. This shows larger impairments in DA release, TTP, and T1/2 in dStr than NAc (multiple paired t-test with Holm-Sidák correction for multiple testing, #p<0.05, ###P<0.001). Data are means±SEM. Data are means±SEM. N= 9 WT slices and 7 DAT-I312F/D421N$^{+/+}$ mice slices.

| | WT | | | DAT-I312F/D421N$^{+/+}$ | | | DAT-I312F/D421N$^{+/+}$ % of corresponding WT region | | |
|---|---|---|---|---|---|---|---|---|---|
| | Striatum total | dStr | NAc | Striatum total | dStr | NAc | Striatum total | dStr | NAc |
| **TTP (s)** | 0.24±0.02 | 0.24 ±0.02 | 0.28±0.02 | 0.66±0.06 **** | 0.67 ± 0.09 ** | 0.61±0.05 *** | 240±30 *** | 280 ± 50 ** | 210±20 ***/# |
| **T1/2 (s)** | 0.42 ±0.02 | 0.42 ±0.03 | 0.46 ±0.02 | 2.2 ±0.2 **** | 2.4 ± 0.3 *** | 2.1 ±0.2 *** | 500 ±50 **** | 580 ± 80 ** | 460 ±50 ***/# |
| **DA peak (µM)** | 1.0±0.05 | 1.00±0.08 | 0.95±0.05 | 0.16±0.02 **** | 0.060±0.02 **** | 0.22±0.03 **** | 16±3 **** | 6.0±2 **** | 23±3 ****/### |
| ***significantly different from corresponding region in WT mice** | | | | | | | | | |
| **#significantly different from dStr** | | | | | | | | | |

**Supplementary table 2. Primary antibodies**

| **Target** | **species** | **supplier** | **Applied concentration** |
|---|---|---|---|
| TH | rabbit | Thermo Scientific (#OPA1-04050) | 1:1000 |
| TH | mouse | Millipore (#MAB318) | 1:1000 |
| DAT | rat | Millipore (#MAB369) | 1:1000 |

| | | | |
|---|---|---|---|
| VMAT2 | rabbit | kindly provided from Gary Miller's Lab | 1:1000 (IHC)<br>1:2000 (WB) |
| DR1 | rat | Sigma Aldrich (#D2944) | 1:500 (IHC)<br>1:1000 (WB) |
| DR2 | rabbit | Synaptic Systems (#376 203) | 1:500 (IHC)<br>1:1000 (WB) |

### SUPPLEMENTARY MOVIES

**Supplementary movies 1-4.** Show example video recordings of WT (movie 1+3) and DAT-I312F/D421N$^{+/+}$ mice (movie 2+4) 30 min (movie 1+2) and 120 min (movie 3+4) into an OFT.

**Supplementary movies 5-6.**
Recordings of clasping behavior during tail suspension shown at 50% speed. Supplementary movie 5 displays a WT mouse and supplementary movie 6 a DAT-I312F/D421N$^{+/+}$ mouse with hindlimb clasping.
